# A Conserved Architecture of Allosteric Communications and Regulatory Hotspots in the KRAS Complexes with Diverse Binding Proteins Controls Mechanisms of Effector Mimicry and Allosteric Modulation : Atomistic Revelations from Molecular Simulations and Mutational Scanning of Binding Energetics and Allostery

**DOI:** 10.1101/2025.05.19.654934

**Authors:** Mohammed Alshahrani, Vedant Parikh, Brandon Foley, Gennady Verkhivker

## Abstract

KRAS is a pivotal oncoprotein that regulates cell proliferation and survival through interactions with downstream effectors such as RAF1. Despite significant advances in understanding KRAS biology, the structural and dynamic mechanisms of KRAS allostery remain unresolved. In this study, we employ microsecond molecular dynamics simulations, mutational scanning and binding free energy calculations together with dynamic network modeling to dissect how engineered DARPin proteins K27, K55, K13, and K1 engage KRAS through diverse molecular mechanisms ranging from effector mimicry to conformational restriction and allosteric modulation. Mutational scanning across all four DARPin systems identifies a core set of evolutionarily constrained residues that function as universal hotspots in KRAS recognition. KRAS residues I36, Y40, M67, and H95 consistently emerge as critical contributors to binding stability. Binding free energy computations show that despite similar binding modes K27 relies heavily on electrostatic contributions from major binding hotspots while K55 exploits a dense hydrophobic cluster enhancing its effector-mimetic signature. Allosteric binders K13 and K19, by contrast, stabilize a KRAS-specific pocket introducing new hinges and bottlenecks that rewire the communication architecture of KRAS without full immobilization. Network-based analysis reveals a strikingly consistent theme: despite their distinct mechanisms of recognition, all systems engage a unifying allosteric architecture that spans multiple functional motifs. This architecture is not only preserved across complexes but also mirrors the intrinsic communication framework of KRAS itself, where specific residues function as central hubs transmitting conformational changes across the protein. By integrating dynamic profiling, energetic mapping, and network-modeling our study provides a multi-scale mechanistic roadmap for targeting KRAS, revealing how engineered proteins can exploit both conserved motifs and isoform-specific features to enable precision modulation of KRAS signaling in oncogenic contexts.

## Introduction

The GTPase KRAS (Kirsten rat sarcoma viral oncogene homolog) is a critical oncogene that is mutated in many human cancers with various sources often reporting different cancer mutation frequency of KRAS. The GTPase KRAS (Kirsten rat sarcoma viral oncogene homolog) is a critical oncogene that is somatically mutated in approximately 10% of all human cancers, with particularly high prevalence in certain malignancies. Specifically, KRAS mutations are found in ∼90% of pancreatic adenocarcinomas, ∼40% of colorectal adenocarcinomas, ∼35% of lung adenocarcinomas, and ∼20% of multiple myeloma cases [1–6]. Among the RAS oncogene family—which includes KRAS, HRAS, and NRAS—KRAS is the most frequently mutated, accounting for ∼30% of all RAS-driven cancers, while HRAS and NRAS mutations are less common, occurring in ∼8% and ∼3% of cases, respectively [7,8]. These mutations drive tumorigenesis by disrupting normal cellular signaling pathways, leading to uncontrolled cell proliferation and survival. RAS proteins function as binary molecular switches, cycling between an active GTP-bound state and an inactive GDP-bound state. This cycling is tightly regulated by two classes of proteins: Guanine nucleotide exchange factors (GEFs), which promote the exchange of GDP for GTP, activating RAS, and GTPase-activating proteins (GAPs), which enhance the intrinsic GTPase activity of RAS, facilitating GTP hydrolysis and returning RAS to its inactive state [9].

A complete GTPase reaction requires well-ordered conformations of the KRAS active site, which includes the phosphate-binding loop, P-loop (residues 10–17), switch I (residues 25–40) and switch II (residues 60–76) regions [10–12] (Figure 1). KRAS can adopt both open and closed states in its active (GTP-bound) form, depending on the conformation of its switch regions (Switch I and Switch II). These states are dynamic and play a critical role in regulating KRAS signaling and interactions with effector proteins. In the GTP-bound state, Switch I and Switch II undergo conformational changes that are critical for effector protein binding. Historically, it has been assumed that the GTP-bound state predominantly adopts a “closed” conformation (state II), which is competent for effector binding and signaling [13] However, recent studies challenged this oversimplified view by demonstrating that even in the GTP-bound state, KRAS can sample multiple conformations including an “open” state (state I) that is not competent for effector binding. Solution-state nuclear magnetic resonance spectroscopy (NMR) studies of RAS bound to the non-hydrolyzable GTP analog, GMPPNP revealed two dominant conformational states in the switch I defined as state I (open conformation) and state II (closed conformation) that exist in the dynamic equilibrium [14–16]. In the state I switch I region adopts an open conformation that is incompatible with effector binding. This conformation was described as “inactive” because it does not support downstream signaling, despite being in the GTP-bound state. In the state II, switch I adopts a closed conformation that facilitates high-affinity binding to effector proteins like RAF1. These NMR studies demonstrated that the GTP-bound state of KRAS is not a single, static conformation but rather a dynamic ensemble of states, some of which are signaling-competent (closed) and others signaling-incompetent (open). Most oncogenic mutations in RAS proteins, particularly at residues G12, G13, and Q61, impair GTP hydrolysis and lock RAS in its active, GTP-bound conformation. These mutations introduce steric hindrance or alter the local structure of RAS, preventing GAPs from binding effectively. For example, mutations such as G12C, G12D, and Q61H alter the conformational dynamics of KRAS, leading to increased signaling activity and resistance to GAPs [17–20]. As a result, RAS proteins remain constitutively active, leading to the overactivation of downstream signaling pathways such as the MAP kinase pathway (RAS-RAF-MEK-ERK) and the PI3K pathway (PI3K-AKT-mTOR). These pathways drive cell proliferation, survival, and metabolism, contributing to tumorigenesis [21].

**Figure 1.**
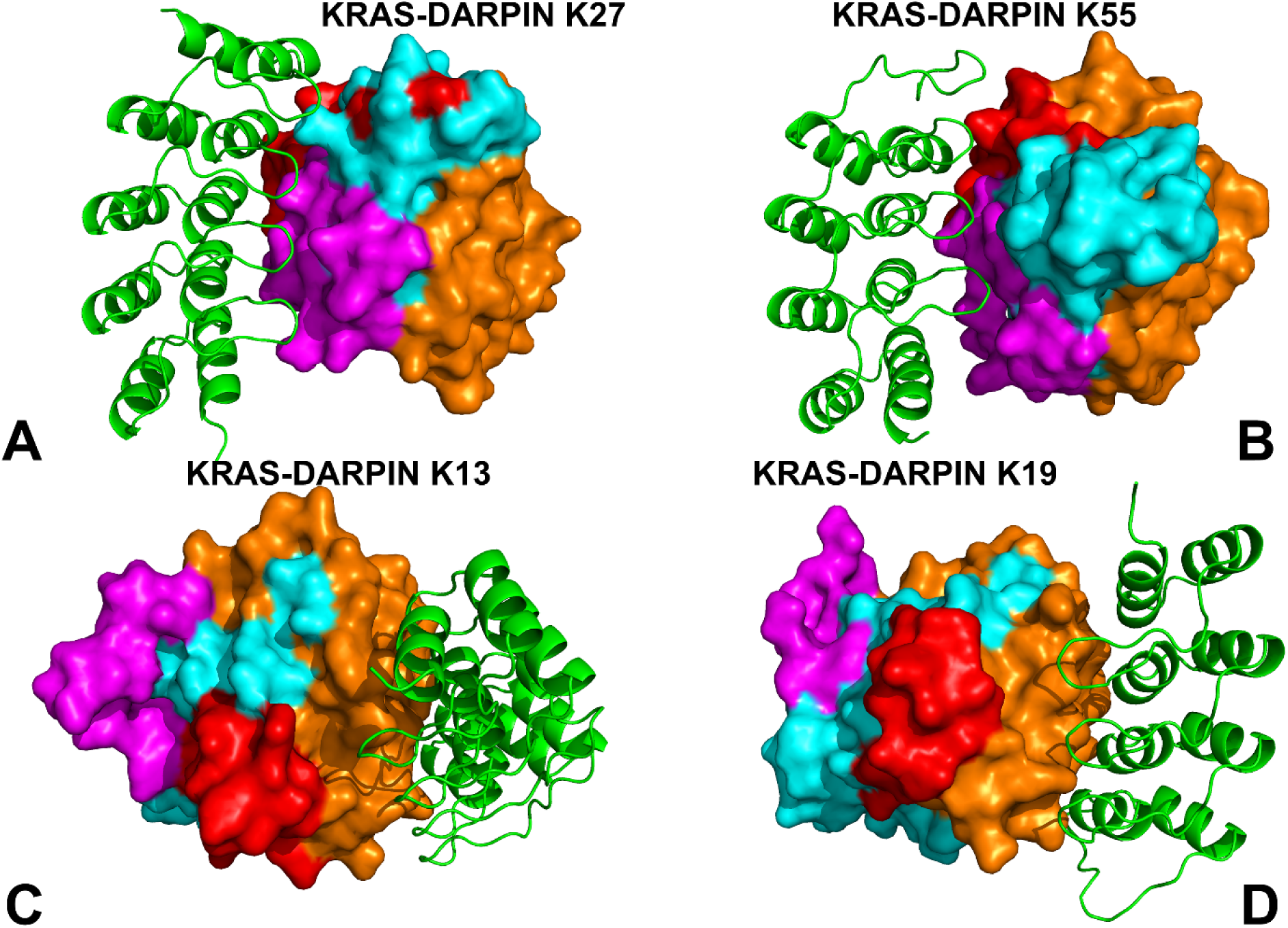
Structural overview and organization of the KRAS protein complexes with DARPin proteins K27 (A), K55 (B), K13 (C) and K19 (D). The crystal structures of KRAS in complexes with DARPin K27 (pdb id 5O2S), DARPin K55 (pdb id 5O2T), DARPin K13 (pdb id 6H46) and DARPin K19 (pdb id 6H47) are used for the analysis and visualization. KRAS protein is shown in surface representation, and DARPin protein binding partners are shown in green ribbons. The functional KRAS regions are highlighted : switch I (residues 24-40 in magenta surface), switch II, (residues 60-76 in red surface), allosteric KRAS lobe (residues 87-166 in orange surface). The remaining KRAS residues are shown in cyan-colored surface representation.

The interaction between KRAS and RAF1 (CRAF) is a critical step in the activation of the MAPK pathway, which regulates cell growth, differentiation, and survival. RAF1 is a serine/threonine kinase that binds to KRAS through two domains: the RAS-binding domain (RBD) and the cysteine-rich domain (CRD). Structural studies have provided detailed insights into the molecular mechanisms underlying this interaction. High-resolution crystal structures of KRAS bound to the RBD of RAF1 have revealed that the RBD interacts primarily with the switch I (residues 25–40) and switch II (residues 60–76) regions of KRAS [22,23]. These regions undergo significant conformational changes during the GTPase cycle, transitioning between disordered (inactive) and ordered (active) states. The N-terminal region of BRAF, including the CR1 and CR2 domains, remains unresolved in the absence of RAS^20^ which suggests that RAS binding is essential for stabilizing these domains and enabling proper RAF activation. Recent cryo-electron microscopy (cryo-EM) studies of full-length BRAF in complex with 14-3-3 and MEK have provided unprecedented views of RAF’s structural biology [24–28]. Oncogenic mutations in KRAS, such as G12V, G13D, and Q61R, do not disrupt the overall interaction with RAF1 but induce local rearrangements in the switch regions. X-ray crystallography of KRAS^Q61H^:GTP revealed that a hyperdynamic switch II allows for a more stable interaction with switch I, suggesting that enhanced RAF activity arises from a combination of absent intrinsic GTP hydrolysis activity and increased affinity for RAF [29]. The nanodisc platforms and paramagnetic relaxation enhancement (PRE) analyses were employed to determine the structure of a hetero-tetrameric complex comprising KRAS and the RBD and CRD of activated RAF1 [30]. By leveraging these advanced techniques, this study investigated how the binding of the RBD or the RBD–CRD differentially modulates the dimerization modes of KRAS on both anionic and neutral membranes. A key discovery of this work is that RBD binding allosterically generates two distinct KRAS dimer interfaces, which exist in a dynamic equilibrium. One interface is favored when KRAS is free, while the other is stabilized when KRAS is in complex with the RBD–CRD [30]. This allosteric regulation highlights the plasticity of KRAS dimerization and its responsiveness to effector binding. Biochemical and structural analyses of variants identified in a KRAS-G12D revealed that attenuation of oncogenic KRAS can be mediated by protein instability and conformational rigidity, resulting in reduced binding affinity to effector proteins or reduced SOS-mediated nucleotide exchange activity. These studies define the landscape of single amino acid alterations that modulate the function of KRAS [31].

Designed Ankyrin Repeat Proteins (DARPins), such as K27 and K55, represent a class of synthetic binding proteins that engage the switch regions of KRAS with high affinity and specificity. These engineered proteins offer promising alternatives to traditional antibodies due to their small size, high stability, and modular recognition capabilities. Engineered protein binders such as DARPins K27 and K55 offer alternative strategies for modulating KRAS function: K27 stabilizes the GDP-bound inactive state, interfering with SOS1-mediated activation, while K55 mimics natural effectors, engaging the active-state conformation and disrupting downstream signaling [32]. These systems—natural regulators and synthetic binders alike—highlight the importance of both conserved structural motifs and flexible interaction networks in shaping KRAS functionality. Another study reported design of allosteric DARPin binders K13 an K19 that specifically inhibit the KRAS isoform by binding to an allosteric site encompassing the region around KRAS-specific residue histidine 95 at the helix α3/loop 7/helix α4 interface [33]. Binding by the DARPin proteins in the allosteric lobe of KRAS influences KRAS-effector interactions modulating KRAS nucleotide exchange and inhibiting KRAS dimerization at the plasma membrane. These results highlighted the importance of targeting the α3/-α4 interface, a previously untargeted site in RAS, for specifically inhibiting KRAS function [33]. A comprehensive analysis of KRAS energetics and allostery was undertaken using a combination of structural biology and biophysical techniques to map the energetic and allosteric landscape of KRAS [34]. This seminal study characterized allosteric energy landscapes that highlighted the thermodynamic consequences of mutations across KRAS providing a quantitative framework to determine allosteric binding sites and their effect on KRAS binding. The study revealed the existence of extensive allosteric networks within KRAS that connect distant regions of the protein. These networks facilitate communication between the nucleotide-binding site, effector-binding site, and other regulatory regions. The authors also identified several cryptic pockets on the surface of KRAS that are transiently formed during conformational changes [34] and can be targeted by small molecules to modulate KRAS activity.

While crystallography and cryo-EM have provided valuable structural insights, computational studies and molecular modeling have played an equally important role in elucidating the dynamics, energetics, and molecular mechanisms of KRAS-RAF1 binding. MD simulations, free energy calculations, and docking studies have complemented experimental techniques, revealing the role of conformational flexibility, membrane interactions, and allosteric regulation. MD simulations have revealed how mutations affect mobility of the switch I and switch II regions of KRAS adopting multiple conformations that influence binding to RAF1 [35,36]. Multiscale coarse-grained and all-atom MD simulations of KRAS4b bound to the RBD and CRD domains of RAF-1, both in solution and anchored to a model plasma membrane explored how RAS membrane orientation relates to the protein dynamics within the RAS-RBDCRD complex [37]. Solution MD simulations describe dynamic KRAS4b-CRD conformations, suggesting that the CRD has sufficient flexibility in this environment to substantially change its binding interface with KRAS4b. Quantitative spatial imaging and atomistic molecular dynamics simulations to examine molecular details of K-Ras plasma membrane binding [38]. The effects of amino acid variations on the structure and dynamics of KRAS-WT and oncogenic mutants G12D, G12V, and G13D of HRAS and KRAS proteins. Based on data from µs-scale molecular dynamics simulations, we show that the overall structure of the proteins remains similar but there are important differences in dynamics and interaction networks [39]. The insightful study employed all-atom molecular dynamics simulations of the C-RAF RBD and CRD regions when bound to oncogenic KRAS4B suggesting that the membrane plays an integral role in regulating the configurational ensemble of the complex that samples a few states dynamically, reflecting a competition between CRAF CRD- and KRAS4B-membrane interactions [40]. Microsecond MD simulations to unveil the binding mechanisms of the FDA-approved MEK inhibitor trametinib with KRASG12D, providing insights for potential drug repurposing. The binding of trametinib was compared with clinical trial drug MRTX1133, which demonstrates exceptional activity against KRASG12D, for better understanding of interaction mechanism of trametinib with KRASG12D [41].

Multiple microsecond MD simulations and Markov State Model (MSM) analysis probed kinetics of MRTX1133 binding to KRAS G12D revealing the kinetically metastable states and potential pathways of MRTX1133 binding while MM/GBSA analysis identified 8 critical residues for MRTX1133 recognition and binding [42]. KRAS binding pockets and ligand interactions, identifying three distinct sites: the conserved nucleotide-binding site, the shallow switch-I/II pocket, and the allosteric Switch-II/α3 pocket [43]. In addition, this study identified KRAS-inhibitor interaction fingerprints aided by MD simulations that characterized the flexibility of these sites to accommodate diverse ligands [43]. MD simulations of the KRASG12C-AMG 510 complex investigated the impact of ligand binding on KRASG12C conformational changes [44]. Mapping simulation trajectories onto the Principal Component Analysis (PCA) model revealed that KRASG12C-AMG 510 predominantly adopted the inactive conformation where AMG 510 reduced the flexibility of the switch regions, stabilizing the KRASG12C-AMG 510 complex in the inactive state [44]. MD simulations explored the structural dynamics and stability of wild-type KRAS and its oncogenic variants (G12C, G12D, G12V, G13D) followed by the free energy landscape analysis revealing unique conformational dynamics and altered thermodynamic stability in mutated KRAS variants [45]. Multiple MD simulations were performed on wild-type and mutant KRAS structures to investigate how G12C and G12D mutations stabilize the active state and how AMG-510 and MRTX1133 inhibitors force these mutants into the inactive state [46]. The study revealed that binding of AMG-510 (Sotorasib) can induce significant stabilization of the Switch-II region of KRAS-G12C, surpassing that of MRTX1133 bound with KRAS-G12D mutant. Mutation probabilities of KRAS-G12 missense mutants and their long-timescale dynamics were assessed by atomistic MD simulations (170 μs) and MSM analysis revealing allosteric hydrophobic signaling network in KRAS, and allosteric modulation of protein dynamics among the G12X mutants which is manifested in the switch regions that are responsible for the effector protein binding [47]. GaMD simulations were performed on the GDP-bound wild-type, G12A, G12D, and G12R KRAS to probe mutation-mediated impacts on conformational alterations of KRAS, showing that all three G12 mutations can alter the structural flexibility of the switch domains [48]. The analyses of the free energy landscapes indicated that the examined G12 mutations can induce more conformational states of KRAS and particularly result in more disordered switch domains [48]. GaMD simulations followed by deep learning (DL) were conducted to probe the effect of G12C mutation and binding of three proteins NF1, RAF1, and SOS1 on the conformational dynamics and free energy landscapes of KRAS4B, showing that mutation and binding alter contacts in key structural domains and enhance switch I and switch II mobility [49]. These findings highlighted the roles of partner binding and G12C in KRAS4B activity and allosteric regulation, providing theoretical insights into KRAS4B function. GaMD simulations and DL also explored the phosphorylation-mediated effect on conformational dynamics of the GTP-bound KRAS revealing that the phosphorylation of pY32, pY64, and pY137 sites leads to more disordered states of the switch domains and can induce conformational transformations between the closed and open states [50]. Overall, structural and computational studies showed that in its active state KRAS predominantly adopts a stable closed conformation that allows for productive interactions with effector proteins and signaling. KRAS can also transiently adopt open and intermediate states, especially during conformational fluctuations or when interacting with inhibitors. The balance between these states is influenced by nucleotide binding, oncogenic mutations, post-translational modifications, and interactions with partner proteins.

Despite the significant body of structural, biochemical and computational studies of KRAS dynamics and binding, there are a number of open questions from understanding the fundamental biophysics of KRAS dynamics and allosteric networks to addressing practical challenges in allosteric drug design and resistance. The important open questions include the following issues : (a) How do long-timescale conformational changes in KRAS, particularly in the switch regions influence its binding to effectors (e.g., RAF1, PI3K, RALGDS) and regulators (e.g., GAPs, GEFs)? (b) What are allosteric networks that connect distal mutation sites to the switch regions, and how do these networks modulate KRAS activity? (c) How do KRAS mutations alter the free energy landscape of conformational states, and what are the thermodynamic drivers of mutation-induced oncogenic activation? (e) What are the energetic contributions of specific residues and interactions to KRAS stability, and allosteric effector binding?

In this study, we attempted to address some of these issues by employing an integrative computational simulation strategy that employed multiple microsecond MD simulations of KRAS-DARPIN complexes, systematic mutational scanning of KRAS residues for binding and stability, rigorous MM-GBSA binding free energy analysis and network modeling of allosteric networks to determine binding affinity hotspots and allosteric binding centers for KRAS complexes with a diverse panel of DARPIN proteins K27, K55, K13 and K19. These simulations provide a detailed understanding of conformational flexibility and allosteric communication within the KRAS-DARPIN interactions and the interplay between allostery, binding and dynamics. The ensemble-based mutational profiling of KRAS residues using knowledge-based energy model enabled accurate identification of binding affinity and protein stability hotspots. The detailed MM-GBSA analysis of binding energetics reveals the thermodynamic drivers of binding and the energetic contributions of the binding affinity hotspots to KRAS stability and effector binding. Using MD-inferred conformational ensembles and dynamics-based network modeling, this study mapped potential allosteric hotspots and allosteric communication pathways in KRAS-DARPIN complexes. Consistent with experimental analysis, our results confirmed that the central β-sheet of KRAS acts as a hub for transmitting allosteric signal between distant functional sites, facilitating allosteric communication between the switch regions and the binding interface. Network-based dynamic modeling of allostery predicted key allosteric hotspots in which mutations or structural perturbations can dramatically affect the fidelity of allosteric communication. The results of this study provide important quantitative insights into principles of allosteric communication and allosteric binding in KRAS that are consistent with and can explain the experimental data. In particular, we find that allosteric communication hotspots are enriched close to the functional switch regions and leverage conformational plasticity in these regions for transmitting allosteric signals, also suggesting that local energetic propagation as the main allosteric mechanism.

## Materials and Methods

### Molecular Dynamics Simulations

The crystal and cryo-EM structures of the KRAS and KRAS complexes are obtained from the Protein Data Bank [51] Multiple independent microsecond MD simulations are performed for the crystal structure of wild-type KRAS in complexes with DARPIN K27 (pdb id 5O2S), DARPIN K55 (pdb id 5O2T), DARPIN K13 (pdb id 6H46) and DARPIN K19 (pdb id 6H47).For simulated structures, hydrogen atoms and missing residues were initially modeled using MODELLER program [52] and subsequently reconstructed and optimized using template-based loop prediction approach ArchPRED [53]. The side chain rotamers were additionally optimized by SCWRL4 tool [54]. Protonation states of ionizable residues were assigned using PROPKA, ensuring histidine residues were in the appropriate tautomeric state [55,56]. The tleap module in AMBER was used to generate the topology (prmtop) and coordinate (inpcrd) files. The system was solvated in a TIP3P water box with a 12 Å buffer and neutralized using Na+ or Cl-ions. The final system was saved for further simulations. Molecular mechanics parameters of proteins were assigned according to the ff14SB force field [57,58]. The protein structures were energy-minimized to remove steric clashes and optimize the structure.

All simulation systems were processed through several stages of structure preparation. First, two stages of minimization were performed: (a) with restraints on the protein backbone to relax solvent and ions, and (b) without restraints to minimize the entire system. The minimization was conducted using the pmemd.cuda module with the following parameters: 1,000 cycles of steepest descent followed by 500 cycles of conjugate gradient minimization, a cutoff of 10 Å for non-bonded interactions, and the ff14SB force field for proteins [58,59]. After minimization, the system is heated from 100K to 300K, over 1 nanosecond of simulation time at a constant volume, with integration time 1 fs. Then we relax system at a constant pressure with protein restraints over 1 ns of simulation time at a constant pressure with restraints set to 1 kcal/mol Å^2. Finally, we relax the system with no restraints for 1 ns of simulation time at a constant pressure. This is followed by 2µs of production period with 20,000 frames saved for analysis. We performed multiple independent 2µs MD simulations under NPT conditions (300 K, 1 bar). The temperature was maintained using the Langevin thermostat, and pressure was controlled with the Monte Carlo barostat. A time step of 2 fs was used, and bonds involving hydrogen atoms were constrained using the SHAKE algorithm. MD trajectories were saved every 100 ps for analysis. The ff14SB force field was used for proteins, and the TIP3P model was used for water.

The CPPTRAJ software in AMBER 18 was used to calculate the root mean squared deviation (RMSD) and root mean squared fluctuation (RMSF) of MD simulation trajectories in which the initial structure was used as the reference [59]. The structures of were visualized using Visual Molecular Dynamics (VMD 1.9.3) [60] and PyMOL (Schrodinger, LLC. 2010. The PyMOL Molecular Graphics System, Version X.X.)

To identify the dynamical coupling of the motions between protein segments, the cross-correlation coefficient (C_ij_) was proposed for measuring the motion correlation between the C_α_ atom pair in residues i and j, which is defined as

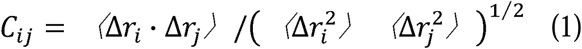

where Δ*r_i_* and Δ*r_i_* are the displacements from the mean position of the C_α_ atom pair in residues *i* and *j* respectively that are evaluated over the sampled period. Positive *C_ij_* are associated with correlated motion in the residue pair, whereas negative *C_ij_* stands for negatively correlated motion. The cross-correlation analysis in this work was implemented using the Bio3D package [61].

### Mutational Scanning and Sensitivity Analysis of the KRAS Residues : Quantifying Effects of Mutations on KRAS Binding and Protein Stability

We conducted a systematic mutational scanning analysis of the KRAS residues in the KRAS complexes using conformational ensembles of KRAS-DARPIN protein complexes and averaging of mutation-induced energy changes. Every KRAS residue was systematically mutated using all substitutions and corresponding protein stability and binding free energy changes were computed with the knowledge-based BeAtMuSiC approach [62–64]. This approach is based on statistical potential describing the pairwise inter-residue distances, backbone torsion angles and solvent accessibilities, and considers the effect of the mutation on the strength of the interactions at the interface and on the overall stability of the complex. The binding free energy of protein-protein complex can be expressed as the difference in the free energy of the complex and free energy of the two protein binding partners:

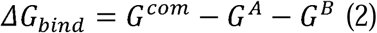

The change of the binding energy due to a mutation was calculated then as the following:

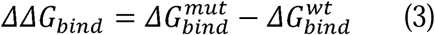

We leveraged rapid calculations based on statistical potentials to compute the ensemble-averaged binding free energy changes using equilibrium samples from simulation trajectories. The binding free energy changes were obtained by averaging the results over 1,000 and 10, 000 equilibrium samples for each of the systems studied.

### MM-GBSA Binding Free Energy Computations of KRAS-DARPIN Complexes

We calculated the ensemble-averaged changes in binding free energy using 1,000 equilibrium samples obtained from simulation trajectories for each system under study. The binding free energies of the KRAS-DARPIN protein complexes were assessed using the MM-GBSA approach [65,66]. The energy decomposition analysis evaluates the contribution of each amino acid to binding of KRAS to DARPIN proteins [67,68]. The binding free energy for the KRAS-DARPIN complexes was obtained using:

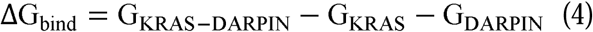

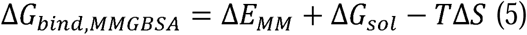

where Δ*E_MM_*is total gas phase energy (sum of Δ*E_internal_*, Δ*E_electrostatic_*, and Δ*Evdw*); Δ*Gsol* is sum of polar (Δ*G_GB_*) and non-polar (Δ*G_SA_*) contributions to solvation. Here, G_KRAS–DARPIN_ represent the average over the snapshots of a single trajectory of the complex, G_KRAS_ and G_DARPIN_ corresponds to the free energy of KRAS and DARPIN protein, respectively. The polar and non-polar contributions to the solvation free energy is calculated using a Generalized Born solvent model and consideration of the solvent accessible surface area. MM-GBSA is employed to predict the binding free energy and decompose the free energy contributions to the binding free energy of a protein–protein complex on per-residue basis. The binding free energy with MM-GBSA was computed by averaging the results of computations over 1,000 samples from the equilibrium ensembles.

First, the computational protocol must be selected between the “single-trajectory” (one trajectory of the complex), or “separate-trajectory” (three separate trajectories of the complex, receptor and ligand). To reduce the noise in the calculations, it is common that each term is evaluated on frames from the trajectory of the bound complex. In this study, we choose the “single-trajectory” protocol, because it is less noisy due to the cancellation of intermolecular energy contributions. This protocol applies to cases where significant structural changes upon binding are not expected. Entropy calculations typically dominate the computational cost of the MM-GBSA estimates. However, for the absolute affinities, the entropy term is needed, owing to the loss of translational and rotational freedom when the ligand binds. In this study, the entropy contribution was not included in the calculations of binding free energies of the complexes because the entropic differences in estimates of relative binding affinities are expected to be small owing to small mutational changes and preservation of the conformational dynamics [69,70]. MM-GBSA energies were evaluated with the MMPBSA.py script in the AmberTools21 package [71].

### Graph-Based Dynamic Network Analysis of KRAS-DARPIN Complexes

To analyze protein structures, we employed a graph-based representation where residues are modeled as network nodes, and non-covalent interactions between residue side-chains define the edges. This approach captures the spatial and functional relationships between residues, providing insights into the protein’s structural and dynamic properties. The graph-based framework allows for the integration of both structural and evolutionary information, enabling a comprehensive analysis of residue interactions. The Residue Interaction Network Generator (RING) program [72–75] was used to generate the initial residue interaction networks from the crystal structures of the KRAS-DARPIN protein complexes. The residue interaction networks for conformational ensembles were constructed by defining edges based on non-covalent interactions between residue side-chains [76,77]. The weights of these edges were determined using two key metrics: (a) dynamic residue cross-correlations derived from MD simulations where these correlations quantify the coordinated motions of residue pairs [78] and (b) coevolutionary couplings measured using mutual information scores, where these couplings reflect evolutionary constraints and residue-residue dependencies [79]. The edge lengths (weights) between nodes ii and jj were computed using generalized correlation coefficients, which integrate both dynamic correlations and coevolutionary mutual information. Only residue pairs observed in at least one independent simulation were included in the network. The matrix of communication distances was constructed using generalized correlations between composite variables that describe both the dynamic positions of residues and their coevolutionary mutual information. This matrix provides a quantitative measure of communication efficiency between residues, reflecting both their physical proximity and evolutionary relationships.

Network graph calculations were performed using the Python package NetworkX [80,81]. This included the computation of key network parameters, such as shortest paths and betweenness centrality, to identify residues critical for communication within the protein structure. The short path betweenness (SPC) of residue *i* is defined to be the sum of the fraction of shortest paths between all pairs of residues that pass through residue *i*:

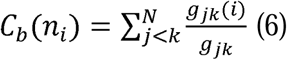

Where *g _jk_* denotes the number of shortest geodesics paths connecting *j* and *k,* and *g _jk_* (*i*) is the number of shortest paths between residues *j* and *k* passing through the node *n_i._* Residues with high occurrence in the shortest paths connecting all residue pairs have a higher betweenness values. For each node *n,* the betweenness value is normalized by the number of node pairs excluding *n* given as(*N* -1)(*N* - 2) / 2, where *N* is the total number of nodes in the connected component that node *n* belongs to.

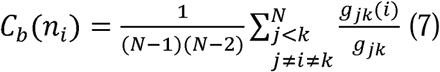

To account for differences in network size, the betweenness centrality of each residue ii was normalized by the number of node pairs excluding ii. The normalized short path betweenness of residue *i* can be expressed as : *g _jk_* is the number of shortest paths between residues *j* and k; *g _jk_* (*i*) is the fraction of these shortest paths that pass through residue *i.*Residues with high normalized betweenness centrality values were identified as key mediators of communication within the protein structure network.

### Network-Based Mutational Profiling of Allosteric Residue Centrality

Through mutation-based perturbations of protein residues we compute dynamic couplings of residues and changes in the short path betweenness centrality (SPC) averaged over all possible modifications in a given position. The change of SPC upon mutational changes of each node is reminiscent to the calculation of residue centralities by systematically removing nodes from the network.

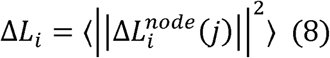

where *i* is a given site, *j* is a mutation and〈⋯〉denotes averaging over mutations. 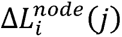 describes the change of SPC parameters upon mutation *j* in a residue node *i*. Δ*L_i_* is the average change of ASPL triggered by mutational changes in position *i*.

Z-score is then calculated for each node as follows:

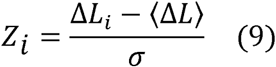

〈Δ*L*〉 is the change of the SPC network parameter under mutational scanning averaged over all protein residues and σ is the corresponding standard deviation. The ensemble-average Z score changes are computed from network analysis of the conformational ensembles of KRAS-DARPIN complexes using 1,000 snapshots of the simulation trajectory.

## Results

### MD Simulations of the KRAS–Protein Complexes Reveal Distinct Dynamic Signatures

To investigate the conformational dynamics and functional flexibility of KRAS in complexes with DARPIN binders we employed crystal structures of bound DARPIN K27 (pdb id 5O2S) K55 (pdb id 52T), K13 (pdb id 6H46) and K19 (pdb id 6H47) (Figure 1). Using these strucrues we conducted microsecond-scale molecular dynamics (MD) simulations of the KRAS– DARPIN complexes. These DARPINs engage KRAS through distinct modes. K27 targets the GDP-bound inactive conformation, acting as a conformational blocker that prevents nucleotide exchange by stabilizing the P-loop and Switch I regions (Figure 1A). K55 mimics natural effectors like RAF1, binding to the GTP-bound active state and engaging both Switch I and II loops (Figure 1B). K13 and K19 recognize a novel allosteric lobe, centered on residues around helix α3–loop–α4, including H95, Y96, L133, S136, and Y137, which are KRAS-specific and not conserved across RAS isoforms (Figure 1C,D).

We used MD simulations of the KRAS–RAF1 complex (PDB ID: 6VJJ) as a representative reference system for understanding how protein–protein interactions modulate KRAS dynamics. MD simulations for KRAS-RAF1 were initiated from the closed conformation of switch I (state II), where GTP is stably bound and coordinated by Mg² ions, T35, and S17. Y32 remains positioned over the nucleotide, stabilizing the active state conformation (Figure 2A). The key structural regions of KRAS are P-loop (residues 10-17), Switch I (residues 25-40) and Switch II regions (residue 60-76) reflecting their functional coupling during nucleotide binding and hydrolysis (Figure 1). The α-helices (α-helix 1: 15–24; α-helix 2: 67–73; α-helix 3: 87–104; α-helix 4: 127–136; α-helix 5: 148–166) and β-sheets (β-strand 1: 3–9; β-strand 2: 38–44; β-strand 5: 109–115; β-strand 6: 139–143) showed moderate correlations with the switch regions, while anti-correlated motions are observed between β-sheets (β-strand 3: 51–57; β-strand 4: 77– 84) and the allosteric lobe (residues 87-166) of KRAS (Figure 1). The root mean square fluctuation (RMSF) analysis provided a detailed view of the residue-wise flexibility of KRAS upon binding to DARPINs K27, K55, K13, and K19 (Figure 2). Using RMSF profiles from all-atom MD simulations, we can identify how each DARPin affects the flexibility of key functional motifs (P-loop, Switch I, Switch II), stabilization of interface regions and differential impact on dynamics between effector-mimicking and allosteric binders. DARPin K27 binds to the GDP-bound inactive conformation of KRAS and functions as a conformational restrainer without inducing large-scale structural changes. In the complex formed between KRAS and DARPin K27, RMSF profiling showed that P-loop (residues 10–17) and Switch I (24–40) had reduced backbone fluctuations, particularly at T35, I36, Y40, and Q70 (Figure 2A). KRAS residues Y32, D33, T35, I36, Y40, and Q70 which form the primary interaction patch are among the most rigid segments of the protein, suggesting that K27 sequesters KRAS in a closed, inactive conformation without inducing global structural changes (Figure 2A). Switch II (residues 60– 70), although engaged to some extent, remains more flexible than Switch I, indicating that K27 does not fully restrict switch motion, but rather modulates dynamic equilibrium toward the GDP-bound state (Figure 2A). The RMSF profile for the KRAS–K55 complex reflect a stronger resemblance to natural effector engagement, with significant immobilization of both Switch I and Switch II loops (Figure 2A). RMSF analysis showed that both Switch I and Switch II become more rigid upon K55 binding, indicating a dual-lock mechanism that enhances effector-like stabilization of the switch regions. Switch II residues Y64 and M67 also show substantial rigidity, indicating that K55 binds to an effector-like orientation, locking these residues into a signaling-ready geometry. The P-loop and upstream β-strands (e.g., β1 and β2) retain moderate flexibility, suggesting that K55 allows some degree of nucleotide sampling (Figure 2A).

**Figure 2.**
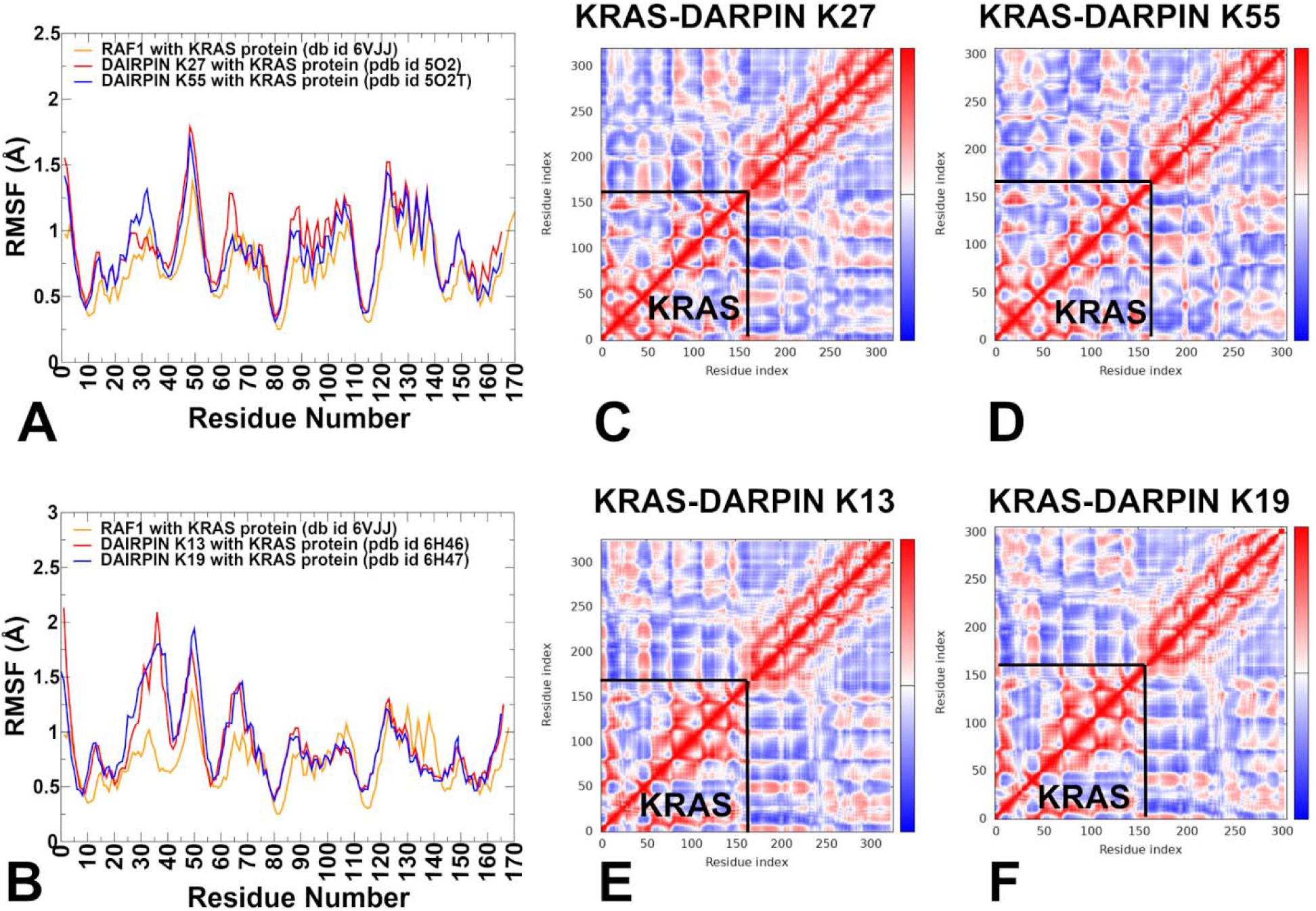
Conformational dynamics of KRAS-DARPin complexes. (A) The RMSF profiles of the KRAS protein residues obtained from MD simulations of the KRAS complexes bound with RBD of RAF1, pdb id 6VJJ (in orange lines), KRAS-K27 complex, pdb id 5OS (in red lines) and KRAS-K55 complex, pdb id 5O2T (in blue lines). (B) The RMSF profiles for the KRAS residues from MD simulations of the crystal structure of wild-type KRAS (bound with the RBD of RAF1, pdb id 6VJJ (in orange lines), KRAS-K13, pdb id 6H46 (in red lines) KRAS-K19, pdb id 6H47 (in blue lines). The DCCM maps for the KRAS residues in the KRAS complex with K27 (C), K55 (D), K13 (E) and K19 (F). DCC measures the correlation between the motions of pairs of residues over the course of MD simulation. Positive correlations indicate that residues move in the same direction, while negative correlations indicate anti-correlated motions. The KRAS protein residues (residues 1-170) are highlighted on the DCCM maps.

Unlike K27 and K55, K13 and K19 bind to a novel allosteric site located at the junction of helix α3–loop–α4, with minimal involvement of the canonical effector interface (Figure 1C,D). This leads to a different pattern of RMSF modulation. For KRAS-K13 complex, allosteric lobe residues H95 and Y96 show reduced mobility, confirming their role as central anchors in this system (Figure 2B). Residues in β-strand 5 and C-terminal loop regions (e.g., L133, S136, Y137) also display lower RMSF values, indicating that K13 stabilizes the structural scaffold supporting the allosteric pocket. Notably, Switch I and Switch II become more mobile, suggesting that K13 does not block functional transitions, but instead alters the conformational ensemble. In KRASB-K19 complex, H95 and R102 showed strong reductions in fluctuation compared to free KRAS or effector-bound KRAS-RAF1 states (Figure 2B). As with K13, Switch I and Switch II loops remain relatively flexible, indicating that K19 modulates function indirectly, possibly by altering signal propagation across the central β-sheet. Hence, K13 and K19 act as dynamic modulators, engaging pockets that are structurally distant from the effector interface, yet capable of influencing long-range communication within KRAS (Figure 2B). The RMSF profiles of KRAS–DARPin complexes clearly distinguish four molecular recognition strategies. K27 locks KRAS into a closed conformation, reducing Switch I dynamics. K55 stabilizes both Switch I and II, resembling natural effectors. K13 and K19 target a KRAS-specific allosteric motif, leading to selective stabilization of the α3–loop–α4 region, while allowing the switch regions to remain partially dynamic.

To understand how different binding partners influence the intrinsic conformational dynamics and residue–residue communication in KRAS, we performed dynamic cross-correlation matrix (DCCM) analysis using MD simulations. This approach maps correlated motions between residues across the protein. The KRAS–K27 complex shows that P-loop and Switch I (residues 10–40) remain tightly coupled, consistent with K27’s preference for the inactive conformation. However, correlations between Switch I and Switch II are reduced, suggesting that K27 may partly suppresses the cooperative motion typically seen in active-state systems (Figure 2C). Positive correlations are reinforced within the P-loop and Switch I. Helix 2 (67–73) and β-strands 5–6 (109–143) —which form part of the central β-sheet —show moderate negative correlations with Switch II, implying that its engagement may indirectly dampen signaling transitions. The KRAS–K55 complex, which mimics natural effectors like RAF1 displays a correlation profile similar to KRAS–K27 with some key distinctions (Figure 2D). Strong positive correlations are maintained between Switch I (25–40) and Switch II (60–76), reflecting K55 ability to stabilize both regions simultaneously. The helix 3 (87–104) and β-strand 4 (77–84) exhibit positive correlations with Switch II, suggesting that K55 binding enhances long-range communication through the central β-sheet. Negative correlations between Switch II and upstream β-strands (e.g., β3 and β4) are diminished indicating that K55 binding stabilizes a more rigid, effector-like network.The KRAS–K13 complex, which engages the allosteric lobe (helix α3–loop–helix α4, residues 87–104), shows a distinct pattern of correlated motion (Figure 2E). Enhanced correlations emerge between the allosteric lobe (residues 87–104) and β-strand 3–4 (51–84), suggesting that K13 modulates signal propagation through the central β-sheet. P-loop and Switch I retain moderate internal correlations, but show reduced coupling with Switch II, indicating that K13 alters the canonical switch–switch communication without fully immobilizing either loop. Residues H95 and Y96, central to K13 binding, become hub residues in the correlation map, showing strong positive correlations with distal regions including Loop 7 (residues 130–140) and helix 5 (residues 148–166), suggesting that K13 introduces new allosteric pathways that extend beyond the effector interface. This pattern indicates that K13 functions not by blocking switch motion, but by modulating the global allosteric network. Similar to K13, K19 targets the allosteric lobe, but with greater involvement of polar interactions (Figure 2F). The H95–D108 motif becomes a central hub, showing strong correlations with both β-strand 4 (77–84) and helix 5 (148–166). Correlation between Switch I and Switch II is preserved, though slightly weakened, indicating that K19 allows for partial switch dynamics. P-loop and helix 1 (15–24) show increased independence from the rest of the structure, suggesting that K19 reduces the global coherence of KRAS motions favoring localized stabilization over broad conformational restriction.

Structral mapping of conformational mobility profiles of KRAS-K27 complex emphasized rigidification of P-loop and Switch I (Supporting Information, Figure S1A). Switch I residues (Y32, D33, T35, I36, Y40) show strongly suppressed mobility, especially around T35 and Y40, which form hydrophobic clusters and hydrogen bonds with the DARPin scaffold. Switch II residues (M67, Q70, Y64) remain moderately flexible, suggesting that K27 does not fully immobilize this loop but instead allows some degree of motion The core β-sheets (β1–β6) remain highly rigid, consistent with their role in maintaining overall fold integrity. The helix α2 and α3 show moderate stabilization, while helix α4 and α5 become slightly more mobile, reflecting allosteric modulation of distal regions due to K27 binding (Supporting Information, Figure S1A). Structural mapping of dynamics profiles for the KRAS–K55 complex highligted greater stabilization of Switch I and Switch II (Supporting Information, Figure S1B). The Y40–M67– F79 aromatic cluster becomes a central rigid domain, showing minimal deviation across simulation time. Structural mapping of dynamic profiles clearly demonstrates that K55 causes more widespread immobilization of KRAS than K27, even in regions far from the immediate binding interface. This is due to several key factors: effector-mimicking engagement of both Switch I and II sand strong hydrophobic and electrostatic interactions centered around Y40, Y64, M67 and R41. In addition, the results suggest enhanced network coupling between functional motifs, including the central β-sheet and helix α5, leading to global rigidification. In contrast, K27 stabilizes the inactive state by focusing on P-loop and Switch I, with minimal impact on Switch II dynamics, resulting in localized stabilization rather than broad conformational restriction. Structural map fo KRAS–K13 complex showed stabilization of the allosteric pocket while poijting to te enhanced switch mobility. K13 acts as a dynamic modulator, where it locally stabilizes the allosteric lobe, but does not restrict global switch motion (Supporting Information, Figure S1C). K19 targets residues H95, Y96, and R102, leading to strong suppression of fluctuations in the allosteric lobe. Structural mapping showed that K19 also induces moderate-to-strong immobilization in regions far from the binding site, such as helix α5 (residues 148–166) and parts of the central β-sheet suggesting that K19 binding alters global signal transmission rather than just local dynamics (Supporting Information, Figure S1D). The widespread immobilization of KRAS regions observed in the presence of K55 and K19, even in areas far from the binding interface, underscores the integrated nature of KRAS dynamics.

### Mutational Scanning of KRAS-DARPin Complexes

Using conformational ensembles obtained from MD simulations, we performed systematic mutational scanning of the KRAS residues in the KRAS-DARPin complexes (Figure 3). In silico mutational scanning was done by averaging the binding free energy changes over the equilibrium ensembles and allows for predictions of the mutation-induced changes of the binding interactions and the stability of the complex. To provide a systematic comparison, we constructed mutational heatmaps for the KRAS interface residues. We used BeatMusic approach to identify binding interface residues. Residues are considered part of the interface if they are within a defined cutoff distance (typically 5 Å) from atoms in the binding partner.

**Figure 3.**
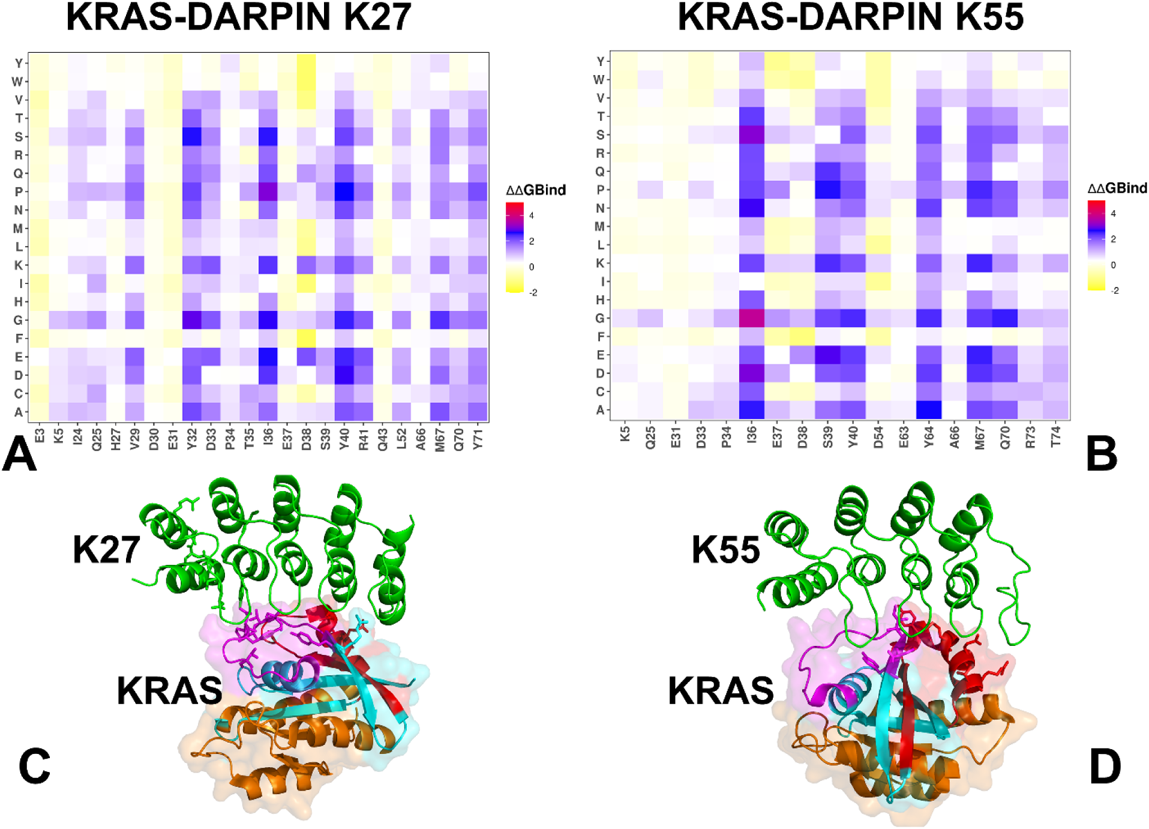
The ensemble-based mutational scanning of binding for the KRAS complexes with DARPin K27 and K55 proteins. The mutational scanning heatmap for the KRAS binding epitope residues in the KRAS complex with DARPin K27 (A) and mutational scanning heatmap for the KRAS binding epitope residues in the KRAS complex with DARPin K55 (B). The binding energy hotspots correspond to residues with high mutational sensitivity. The heatmaps show the computed binding free energy changes for 20 single mutations on the sites of variants. The squares on the heatmap are colored using a 3-colored scale yellow-white-blue-red, with blue color indicating the appreciable unfavorable effects on stability and red pointing to very large destabilizing effects of mutations. The horizontal axis represents the binding epitope residues. Residues are considered part of the interface if they are within a defined cutoff distance 5 Å from atoms in the binding partner. The Y axis depicts all possible substitutions of a given binding epitope residue denoting mutations to letters using a single letter annotation of the amino acid residues. Structural mapping of the binding energy hotspots in the KRAS complexes with DARPin K27 (C) and DARPin K55 (D). The binding epitope hotspot positions for K27 (Q25, V29, Y32, D33, T35, I36, S39, Y40, R41. M67, Q70, Y71) and hotspots for K55 (I36, S39, Y40, Y64, M67, Q70, R73, T74) are shown in sticks. Switch I region (residue 24-40) is depicted in magenta and Switch II region (residues 60-76) is in red. The allosteric lobe (residues 87-166) is shown in orange. The DARPin molecules are shown in green ribbons.

In the KRAS–K27 complex, binding hotspot residues include Y32, D33, T35, I36, Y40, R41, M67, Q70, and Y71 which contribute to its modular, backbone-mediated binding strategy (Figure 3A,C). These residues are primarily located in the Switch I and II regions forming a modular hydrogen bond network that stabilizes the GDP-bound inactive conformation of KRAS. K27 uses a broad, modular recognition strategy where multiple weakly interacting residues collectively stabilize the complex. This enhances its mutation tolerance and makes it particularly effective against variants that stabilize the GDP-bound form. The most critical residue is R41, which forms strong electrostatic interactions with conserved acidic motifs on the DARPin. Substitutions at this site (e.g., R41C/H/N) severely impair binding (Figure 3A,C). Extremely sensitive to mutations affecting T35 and Y40, which play central roles in hydrophobic stabilization and nucleotide coordination.

KRAS shows some resilience to conservative mutations, especially those retaining aromatic character (e.g., Y40F/H) (Figure 3A,C). K27 binding is somewhat less affected by mutations at Switch II residues (e.g., Y64, M67), indicating that it does not rely heavily on this region (Figure 3A,C). Binding of KRAS with DARPin K55 suggests mechanism of effector mimicry through hydrophobic anchoring and electrostatic stabilization. K55 engages both Switch I and Switch II regions, forming a tight hydrophobic core centered around Y40, M67, and F79, while also establishing electrostatic networks involving D38 and R41. The key binding hotspots include I36, S39, Y40, Y64, M67, R41, D38 (Figure B,D) where positions I36, S39, Y40, Y64 and M67 emerged as the least mutation-tolerant. Structural mapping shows that K55 recognizes conserved motifs in Switch I/II, making it ideal for blocking signal propagation rather than nucleotide cycling (Figure 3D). Mutational scanning revealed sensitive to mutations positions that disrupt aromatic stacking, such as Y64G and M67L, which lead to significant destabilization. T35A/N/V substitutions have minimal impact on K55 binding while E37 and D38 substitutions moderately affect binding, reflecting their role in interfacial hydrogen bonding and salt bridge formation (Figure 3B).

However, T35 mutations severely disrupt K27 binding highlighting differences in epitope dependency. Substitutions at Y40 and M67 affect both DARPins, consistent with their central role in hydrophobic stabilization. Similarly, Y64 mutations strongly impair K55 binding due to its reliance on aromatic stacking and side-chain interactions, whereas K27 remains largely unaffected, likely due to its focus on upstream P-loop and Switch I residues. This analysis reinforced the notion that K55 mimics natural effectors like RAF1, relying on localized, high-affinity contacts within the switch regions, which makes it highly potent but also more vulnerable to mutations that affect aromatic or charged residue.

Despite targeting partially overlapping regions on the KRAS surface, only a subset of residues contributes to both K27 and K55 binding. These residues I36, S39, Y40, M67 emerge as shared anchors between the two DARPins, playing roles in stabilizing the interaction. These findings align with experimental observations [32] showing that engineered protein binders can achieve inhibition through complementary strategies.

Both DARPins K13 and K19 engage a novel allosteric lobe on KRAS, located in the α3–loop– α4 motif distinct from the classical effector interface. Despite this shared location, they differ in their residue-specific contributions and polar vs hydrophobic dependency. For KRAS–K13 interactions the binding hotspots are K88, F90, H94, H95, Y96 R102, L133, S136, Y137, G138 (Figure 4A,C) This set of residues forms a central hub in the allosteric lobe, where H95 and Y96 contribute the most to binding via π-stacking and hydrophobic packing with tryptophan residues in the DARPins (W35 and W37) (Figure 4C). In the KRAS-K19 complex, the binding hotspots are F90, H95, E107, D108, L133, S136, Y137 (Figure 4B). K19 shares many key residues with K13 but introduces additional polar interaction networks, particularly at D108 (Figure 4B,D). Mutational scanning confirms that hotspot substitutions severely impair binding in both systems. H94D/E/N and H95D/E/P/V/G lead to strong destabilization in both complexes due to loss of aromatic stacking. R102K/R/D substitutions have stronger impact on K13 where this residue contributes more directly to polar anchoring. E98 mutations (E98A/D/K/P) cause significant destabilization in K19 confirming its importance in maintaining interface compatibility in this system. Conservative mutations such as S136T/A, Y137F/H, and L133V/I are generally better tolerated, supporting their structural compatibility rather than direct functional inhibition.

**Figure 4.**
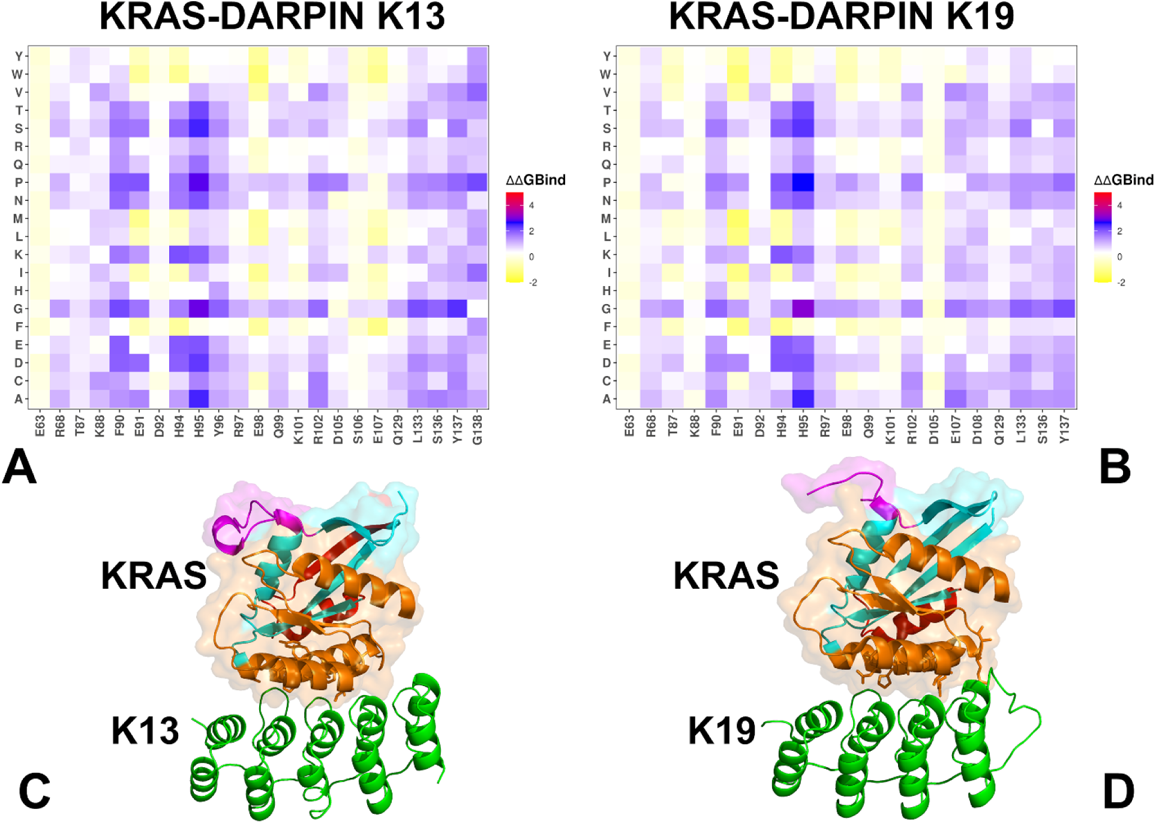
The ensemble-based mutational scanning of binding for the KRAS complexes with DARPin K13 and K19 proteins. The mutational scanning heatmap for the KRAS binding epitope residues in the KRAS complex with DARPin K13 (A) and mutational scanning heatmap for the KRAS binding epitope residues in the KRAS complex with DARPin K19 (B). The binding energy hotspots correspond to residues with high mutational sensitivity. The heatmaps show the computed binding free energy changes for 20 single mutations on the sites of variants. The squares on the heatmap are colored using a 3-colored scale yellow-white-blue-red, with blue color indicating the appreciable unfavorable effects on stability and red pointing to very large destabilizing effects of mutations. The horizontal axis represents the binding epitope residues. Residues are considered part of the interface if they are within a defined cutoff distance 5 Å from atoms in the binding partner. The Y axis depicts all possible substitutions of a given binding epitope residue denoting mutations to letters using a single letter annotation of the amino acid residues. Structural mapping of the binding energy hotspots in the KRAS complexes with DARPin K13 (C) and DARPin K19 (D). The binding epitope hotspot positions for K13 (K88, F90, H94, H95, R102, L133, S136, Y138, G138) and hotspots for K19 (F90, H94, H95, E107, D108, L133, S136, Y137) are shown in sticks. Switch I region (residue 24-40) is depicted in magenta and Switch II region (residues 60-76) is in red. The allosteric lobe (residues 87-166) is shown in orange. The DARPin molecules are shown in green ribbons.

Several KRAS residues are engaged by multiple DARPINs, albeit to different degrees. I36 is important for hydrophobic core integrity in K27 and K55. Y40 is a dominant anchor in K27 and K55, contributing to both hydrophobic and electrostatic interactions. R41 is central residue in K27 and K55, though K27 shows greater reliance on its side chain. H95 is engaged exclusively by K13 and K19, reinforcing their KRAS-specificity. These findings indicate that certain residues, particularly I36 and Y40, are evolutionarily constrained interaction hubs making them ideal targets for broad-spectrum inhibition strategies. The analysis reveals how each binder engages KRAS through distinct mechanisms with both shared and unique energetic hotspots. The presence of unique hotspots suggests that combining these DARPINs could raise the genetic barrier to resistance, since escape would require simultaneous mutation of structurally distant and functionally divergent residues.

These findings highlight that mutational scanning is highly effective in identifying major binding hotspots and predicting the impact of loss-of-function mutations. However, they also reveal limitations in capturing subtle specificity shifts, particularly for polar and charged residues at the periphery of the binding interface. The resolution of the simplified knowledge-based energy function used in our analysis appears sufficient for detecting large-effect mutations (e.g., those involving hydrophobic collapse or charge reversal), but may be somewhat less sensitive to fine-grained changes in electrostatic networks or hydrogen bond rearrangements that may influence binding selectivity. Importantly, the observed conservation of key hotspot residues across natural effectors and engineered binders underscores the presence of conservative mechanisms in KRAS recognition.

### MM-GBSA Analysis of the Binding Energetics Provides Quantitative Characterization of Thermodynamic Drivers of DARPin Interactions

Using the conformational equilibrium ensembles obtained MD simulations of the KRAS-DARPin complexes we computed the binding free energies for these complexes and analyzed the residue-specific contributions to binding (Figures 5,6). The MM-GBSA residue-based decomposition reveals that R41 is the dominant hotspot in the KRAS–K27 interface, contributing ΔG = -16.34 kcal/mol, primarily through electrostatic interactions (ΔGelec = - 137.18 kcal/mol) (Figure 5A). This residue acts as a central anchor, more so than any other in the system. Other notable contributors include Y32, T35, I36, Y40, M67, Q70, Y71 — all showing favorable van der Waals contributions, reinforcing the hydrophobic nature of the K27 interface (Figure 5B). D33 and D38 contribute via polar contacts and hydrogen bonds, playing supportive roles in maintaining overall interface compatibility (Figure 5C). K55 derives its strongest stabilization from van der Waals interactions and engages a tighter, more localized interface with strong contributions from M67 (ΔG_vdw = -4.07 kcal/mol), Y64 (ΔG_vdw = -3.3 kcal/mol), Y40 (ΔG_vdw = -3.45 kcal/mol), I36 (ΔG_vdw = -2.48 kcal/mol) (Figure 5D,E). These residues form a central hydrophobic cluster, especially involving Y40–M67–F79 that mimics natural effectors. This difference highlights how K27 uses a modular, distributed energy model, while K55 exploits a concentrated, high-affinity hotspot network. Electrostatic contributions are also significant, particularly at D33 (ΔG_elec = -50.42 kcal/mol) that forms long-range salt bridges with R41 and R73 in K55 (Figure 5F). E37 and D54 also contribute to polar networks that stabilize Switch I geometry (Figure 5F). Unlike K27, K55 does not rely heavily on R41, though this residue still plays a supportive role in shaping the electrostatic landscape. Both K27 and K55 exploit hydrophobic interactions to stabilize the complex. Y40 contributes significantly via aromatic stacking and I36 and M67 provide favorable van der Waals contributions, forming part of a hydrophobic core that anchors the interface (Figure 5B,E). This reflects the broader theme seen across multiple effectors and inhibitors: hydrophobic clusters play a dominant role in maintaining interface compatibility especially when they involve conserved aromatic or aliphatic side chains. The MM-GBSA results suggested that the electrostatic contributions support the interface integrity (Figure 5C,F). While not the primary driver in all cases, electrostatic interactions contribute to binding stability in both systems. In K27, key contributors include R41, D33, and D38, where long-range charge complementarity enhances binding affinity (Figure 5C). In K55, residues D33, E37, and D54 participate in salt bridges and hydrogen bonds with basic motifs in the DARPin, reinforcing interface compatibility (Figure 5F). Despite differences in how each binder utilizes these interactions, electrostatic networks remain essential for stabilizing the switch regions.

**Figure 5.**
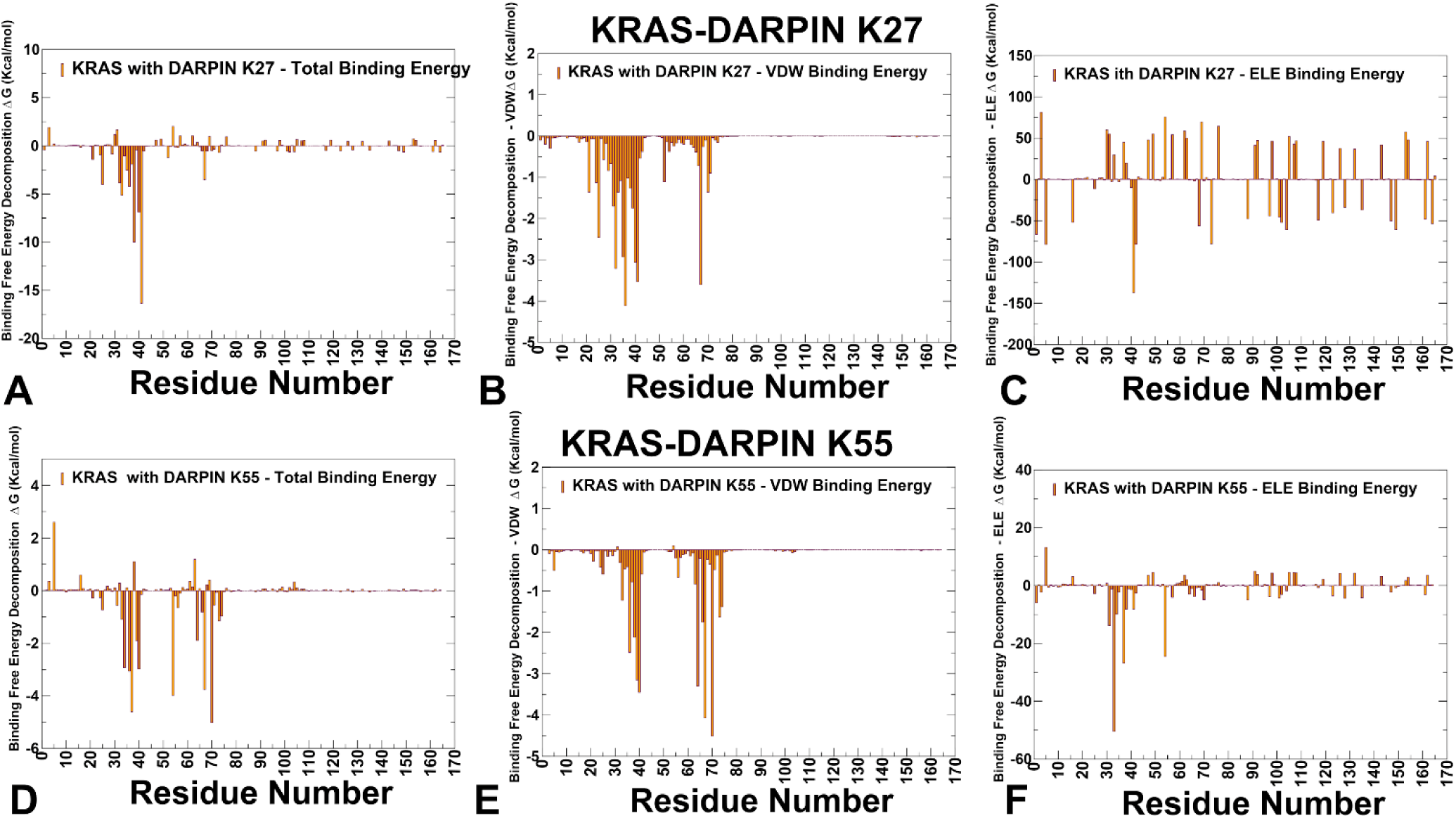
The residue-based decomposition of the total binding MM-GBSA energies for the KRAS residues in the KRAS complex with DARPin K27 (A-C) and DARPin K55 (D-F). The residue-based decomposition of the total binding energy (A), van der Waals contribution (B) and electrostatic contribution to the total MM-GBSA binding energy (C) for the KRAS residues in the KRAS complex with K27. The residue-based decomposition of the total binding energy (D), van der Waals contribution (E) and electrostatic contribution to the total MM-GBSA binding energy (F) for the KRAS residues in the KRAS complex with K55. The MM-GBSA contributions are evaluated using 1,000 samples from the equilibrium MD simulations of respective KRAS complexes.

**Figure 6.**
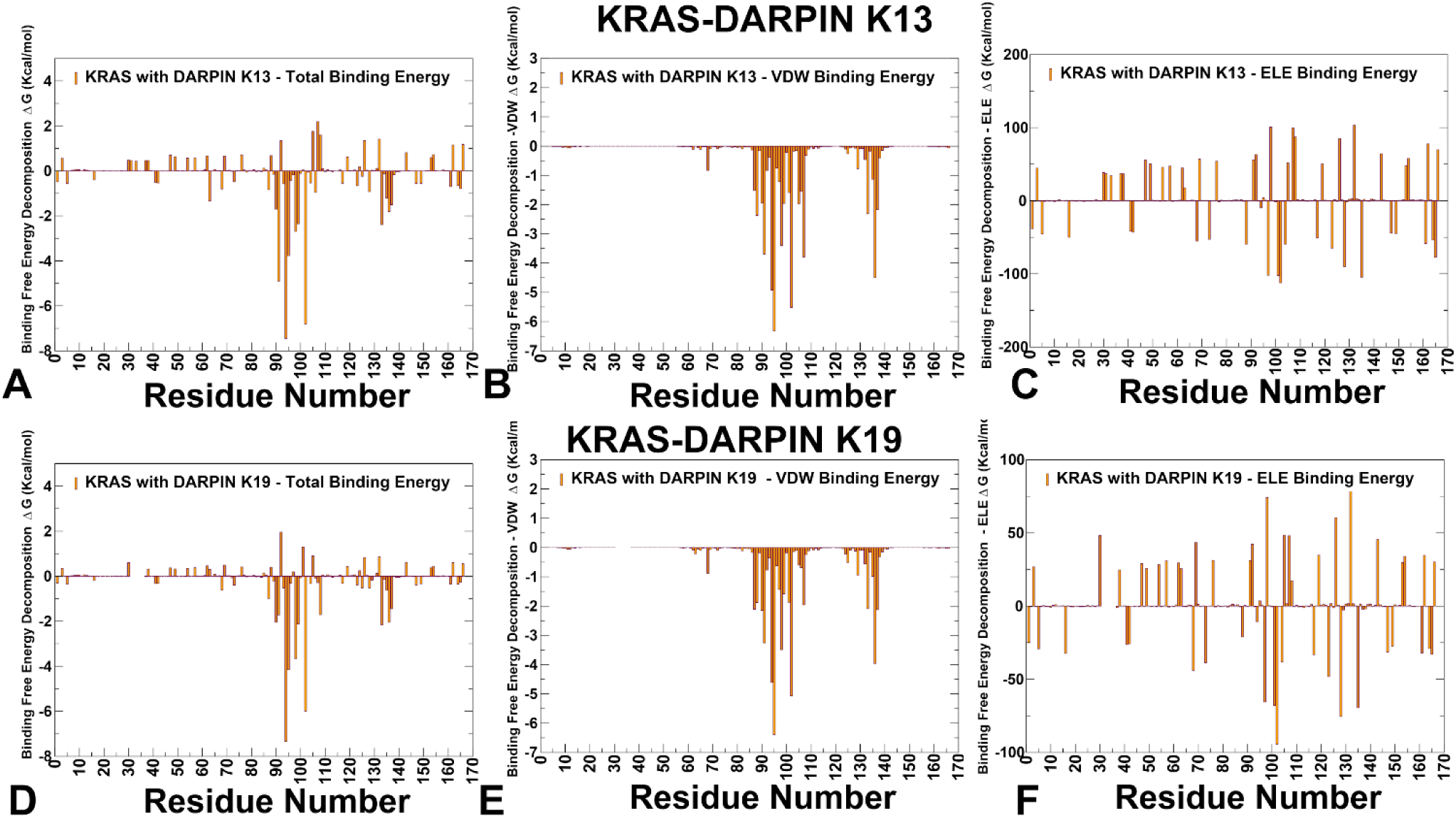
The residue-based decomposition of the total binding MM-GBSA energies for the KRAS residues in the KRAS complex with DARPin K13 (A-C) and DARPin K19 (D-F). The residue-based decomposition of the total binding energy (A), van der Waals contribution (B) and electrostatic contribution to the total binding energy (C) for the KRAS residues in the complex with K13. The residue-based decomposition of the total binding energy (D), van der Waals contribution (E) and electrostatic contribution to the total MM-GBSA binding energy (F) for the KRAS residues in the KRAS complex with K19. The MM-GBSA contributions are evaluated using 1,000 samples from the equilibrium MD simulations of respective KRAS complexes.

The MM-GBSA results clearly distinguish K27 and K55 as two functionally divergent classes of engineered binders. K27 emphasizes electrostatic anchoring and modular hydrogen bonding, stabilizing the GDP-bound inactive conformation. K55 exploits hydrophobic clusters and effector-like electrostatic networks, locking KRAS into a signaling-ready state that blocks downstream effector access. Despite targeting partially overlapping surfaces, these DARPINs differ significantly in their energetic dependencies and sensitivity to mutations. MM-GBSA computations suggested that K27 binding is driven by a dominant electrostatic hotspot at R41, which anchors the interface through salt bridge formation. Hydrophobic contributions from Y32, T35, I36, Y40, M67, Q70, and Y71, forming a modular hydrogen bond network that sequesters KRAS in an inactivation-prone orientation. We argue that the mechanism of K27 binding is determined by modular hydrophobic and electrostatic anchoring. K55 binding is primarily driven by a central hydrophobic core involving Y40, M67, and F79, reinforced by electrostatic networks at D38 and S39. As a results, K55 binding mechanism resembles effector mimicry through compact hydrophobic clustering. These findings support the concept of multi-binder strategies, where combinations of conformational blockers (K27) and effector mimics (K55) can synergistically reduce oncogenic KRAS activity.

DARPins K13 and K19 bind to an evolutionary divergent but structurally conserved allosteric pocket involving H94, H95, Y96, L133, S136 and Y137 (Figure 6). These residues form part of a hydrophobic core that supports structural compatibility between KRAS and the DARPIN scaffold, even though the pocket is not involved in effector recognition or nucleotide exchange. Both systems exploit conserved hydrophobic motifs —especially involving H95 and Y96 — which serve as central anchors for stabilizing the interaction. In both complexes, van der Waals interactions dominate the binding mechanism as H95 contributes significantly through aromatic stacking with tryptophan residues in the DARPin scaffold. L133 and Y137 reinforce shape complementarity, reducing conformational freedom in Loop 7 and helix α5. These findings align with experimental observations showing that mutations affecting His95A/V/N severely impair binding for both K13 and K19, confirming its role as a KRAS-specific hotspot.

Despite their shared allosteric engagement, K13 and K19 differ significantly in their electrostatic contributions and mutation sensitivity, leading to distinct mechanisms of stabilization and isoform selectivity. K13 relies heavily on H94 and R102, forming a dense hydrophobic cluster that stabilizes the α3–loop–α4 interface. It engages E91 and L133 via moderate electrostatic and van der Walls forces. The top contributors to binding stability are H94 > R102 > E91 > H95 (Figure 6A-C). This ranking highlights that K13 relies strongly on E91, which forms polar interactions and hydrogen bonds that support structural compatibility between the DARPin and KRAS. The analysis showed that K13 forms a dense hydrophobic core involving H94 and H95, supported by polar networks formed with E91 and R102 (Figure 6B). The electrostatic contribution of R102 also plays a critical anchoring role (Figure 6C). In contrast, K19 shows a different residue dependency, with the following hotspot ranking H94 > R102 > H95 > E98 (Figure 6D-F). This shift reflects the role of E98, which engages in backbone hydrogen bonding and water-mediated interactions that contribute significantly to interface stability. While E98 does not form direct side-chain contacts with the DARPin, its presence helps maintain local structure and hydration compatibility, enhancing overall binding fidelity. Additionally, H94 shows improved electrostatic compatibility in K19 (Figure 6F), indicating that it participates in enhanced hydrogen bond formation compared to K13. The MM-GBSA analysis of the KRAS– K13 and KRAS–K19 complexes reveals that both DARPINs engage a structurally stable allosteric lobe but do so with different energetic strategies. K13 emphasizes E91 and R102, forming moderate electrostatic networks that support interface integrity. K19 leverages E98 and H94, introducing additional polar constraints that enhance KRAS specificity. K19 shares many of the same hydrophobic anchors as K13 especially at H94 and H95 but introduces new polar constraints that increase KRAS selectivity. In particular, D108 emerged as an additional hotpot for K19 and E98 appeared to be a significant contributor to binding stability. This enhanced electrostatic dependency allows K19 to fine-tune the signaling output of KRAS. Together, these findings highlight how engineered proteins can exploit non-canonical pockets to achieve precision-based targeting, offering complementary advantages when used in combination with effector-mimicking agents such as K55 or RAF1 RBD mimics.

### Network Analysis of Allosteric Communication in KRAS–DARPin Complexes

We used the ensemble-based network centrality analysis and the network-based mutational profiling of allosteric residue propensities to characterize global network of allosteric communications. We constructed residue interaction graphs using the ensemble-averaged contact maps from MD simulations of KRAS complexes with DARPINs K27, K55, K13, and K19. Each node represents a KRAS residue, and edges are defined based on proximity (≤ 5 Å) in the conformational ensemble. The Shortest Path Betweenness (SPC) centrality was computed for each residue, measuring how often it lies on the shortest paths connecting distant parts of the protein. High-SPC residues function as central routers in the network, playing critical roles in information transfer even if they are not directly involved in binding. To assess how mutations perturb this network, we calculated the Z-score of SPC changes across an ensemble of mutation-induced structures. A high positive Z-score indicates that a mutation disrupts communication pathways, suggesting that the wild-type residue contributes significantly to network integrity and signaling robustness Allosteric hotspots are identified as residues in which mutations incur significant perturbations of the global residue interaction network that disrupt the network connectivity and cause a significant impairment of global network communications and compromise signaling.

Network analysis of KRAS-K27 revealed a number of high SPC residues including positions in the interacting KRAS regions (Y40, R41, D38, S39, Y71, V14, K16, L23, I24, F78, F82, F90) as well as residues in the allosteric lobe (L113, S136, L159) (Figure 7A). In the KRAS–K27 complex, SPC analysis also identified several clusters of high-centrality residues: P-loop (residues 10–17) : L6, G15, S17, I18; Switch I (residues 24–40) : Y32, P34, T35, I36, E37, D38, S39; α-helix 1 (residues 80–85) : M80, A83; α-helix 3 (residues 87–104) : R97, H95, Y96; and β-strand 4 (residues 77–84) : L77, V81, L83. Y40 and R41 emerged as the dominant hubs showing the highest SPC value. This residue acts as a central router in the network, mediating long-range communication between the P-loop and C-terminal helices. T35 and Y40 also showed elevated SPC, reinforcing their role in maintaining nucleotide coordination geometry. The central β-sheet, especially β-strand 2 (E37–D38–S39) and β-strand 3 (I55–L56–D57), exhibited increased network centrality, indicating that K27 indirectly stabilizes these motifs, even though it does not engage them directly (Figure 7A). K55 interface includes key aromatic anchors such as Y40, M67, and F79, which form a hydrophobic cluster that locks the switch loops into a signaling-ready configuration. The highest SPC positions in the network centrality profile for KRAS-K55 complex include residues Q70, Y71, Y64, M67, E37, D38, T74, F78, F82, I93, V114, N116, L133, Y137 (Figure 7B). Although the overall shape and patterns of the SPC profiles for K27 and K55 complexes are generally similar as may be expected, there are noticeable differences in relative ranking of the top SPC residues. For KRAS-K55 complex, the SPC profile highlighted the importance of Switch II residues Q70, Y71, Y64, M67 which form a hydrophobic cluster that locks the switch loops into a signaling-ready configuration. The network mediating role of these Switch II residues reflected their critical roles in effector mimicry and hydrophobic stabilization (Figure 7B). E37 and D38, known for their involvement in salt bridge formation with RAF1 also showed high SPC values, suggesting they could function as local hubs in the effector lobe. Importantly, among SPC profile peaks are residues F78, L80, and F82 from β-strand 4; I93 from α-helix 3; V114 and N116 from β-strand 5 and Y157, L159, I163 and R164 from α-helix 5 of the allosteric lobe (Figure 7A,B).

**Figure 7.**
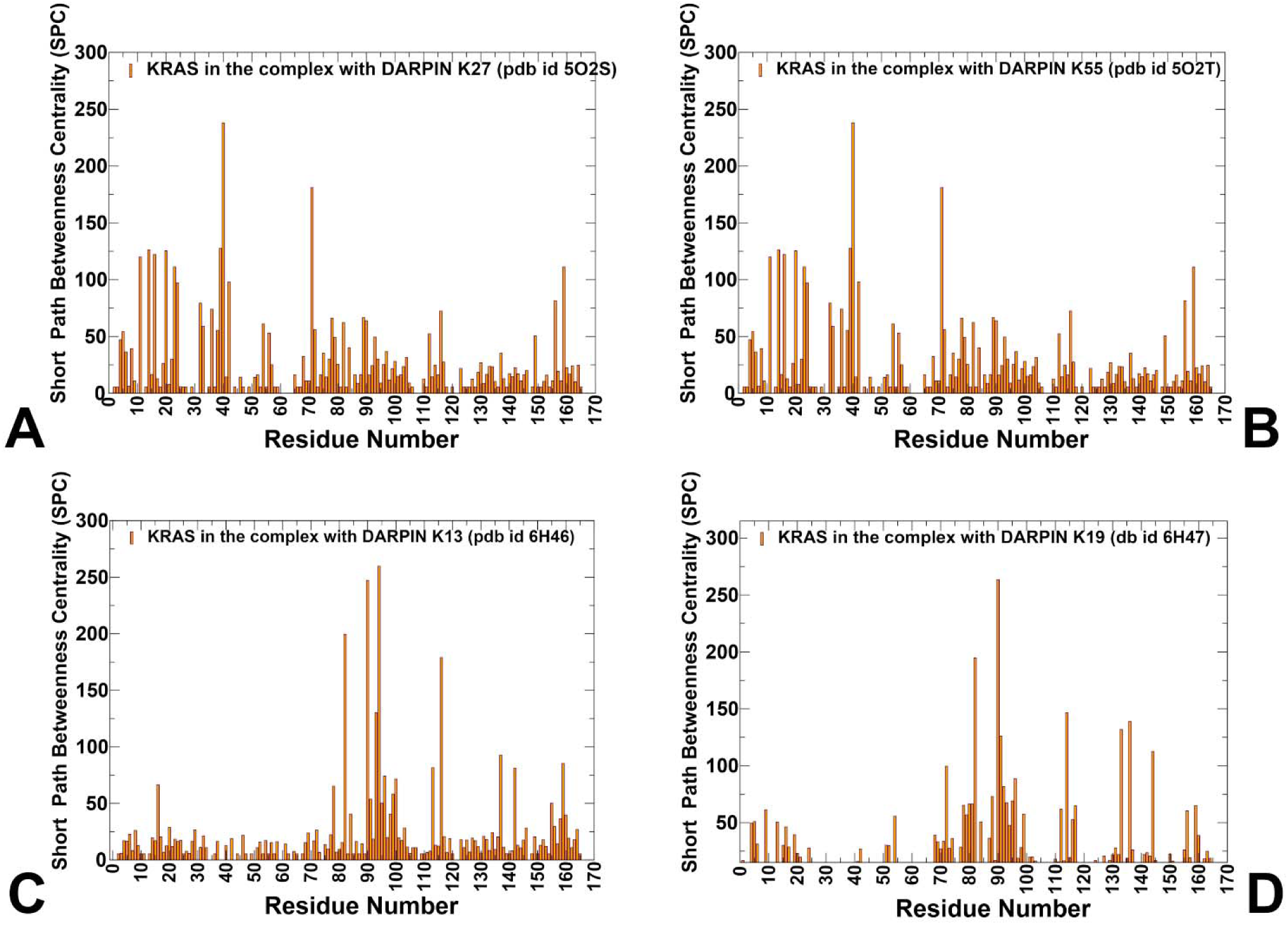
Network centrality analysis of the KRAS complexes with DARPIN proteins. The ensemble-averaged SPC centrality for the KRAS residues in the KRAS-WT complex with DARPIN K27 (pdb id 5O2S) (A), KRAS-WT complex with DARPIN K55 (B), KRAS-WT complex with DARPIN K13 (C) and KRAS-WT complex with DARPIN K19 (D). The profiles are shown in orange-colored filled bars.

Hence, the SPC analysis highlighted the presence of allosteric mediating centers from different functional regions including positions located away from the binding interface. The results also highlight the importance of allosteric communication in KRAS, where distant residues influence the binding interface with RAF1 through long-range interactions. This communication is mediated by a network of residues that function as bridges or bottlenecks in the allosteric network.

Unlike K27 and K55, which target the effector lobe, K13 and K19 bind to a novel allosteric pocket involving helix 3–loop–helix 4, centered around H95, Y96 and surrounding motifs. SPC analysis revealed elevated centrality values for F78, H94, F90, F82, N116, I93, Y137, L159, L113, I142, H95, Y96 (Figure 7C). H95, Y96 form the core of the allosteric hub while L133, S136, Y137 provide allosteric communication centers For KRAS-K19, the major allosteric centers are F82, F90, V114, S136, E91, T144, M72, H95, Y96 (Figure 7D). These residues do not show strong overlap with those identified in effector systems, suggesting that K13 and K19 engage a structurally and dynamically distinct region of the network, offering a modular route for isoform-specific targeting. Importantly, all KRAS-DARPin complexes share a number of common allosteric hotspots including F78, F82 (β-strand 4) F90, I93 (α-helix 3),V114, N116 (β-strand 5), that are located in different functional regions of KRAS (Figure 7, Supporting Information, Figure 2). Structural mapping showed that many of these residues form communication routes via the central β-sheet, linking distal pockets to the active-site lobe (Supporting Information, Figure S2). The predicted allosteric hotspots are in strong agreement with experimental data [34], which independently identified residues in β-strand 4 (F78, F82) and β-strand 5 (V112, V114, N116) as critical mediators of long-range communication, linking distant functional sites and stabilizing specific conformations. These hotspots, which align strongly with experimental data, underscore a conserved allosteric infrastructure that is universally important for KRAS signaling. The findings highlight a conserved allosteric infrastructure that can be exploited for therapeutic design.

By comparing the top-ranked residues based on their Z-scores, we can gain deeper insight into the distinct mechanisms by which these DARPINs engage KRAS. While mutational scanning identified hotspot residues such as R41, Y40, T35, I36, and M67 as major contributors to interface stability, the Z-score SPC analysis revealed additional mutation-sensitive control points that are not part of the direct DARPin interface yet have profound effects on network integrity. Network-based mutational scanning revealed that certain residues—although not part of the direct interface—strongly affect binding when mutated. Among top Z-score profile residues are Y40, Y71, T20, K16, V14, L23, F78, S39, N116 (Figure 8A). These findings suggest that K27’s mechanism extends beyond localized stabilization. For K55, the top Z-score SPC residues identified are quite different and include Q70, Y71, T74 along with residues in the allosteric lobe Y137, L133, R97, A134. (Figure 8B). These residues highlight regions that are central to signal propagation and network stability, even though they may not directly interact with the DARPin. Mutations at these network-embedded residues even if they do not contact the DARPin directly, can severely disrupt information flow leading to reduced binding affinity or altered functional outcomes.

**Figure 8.**
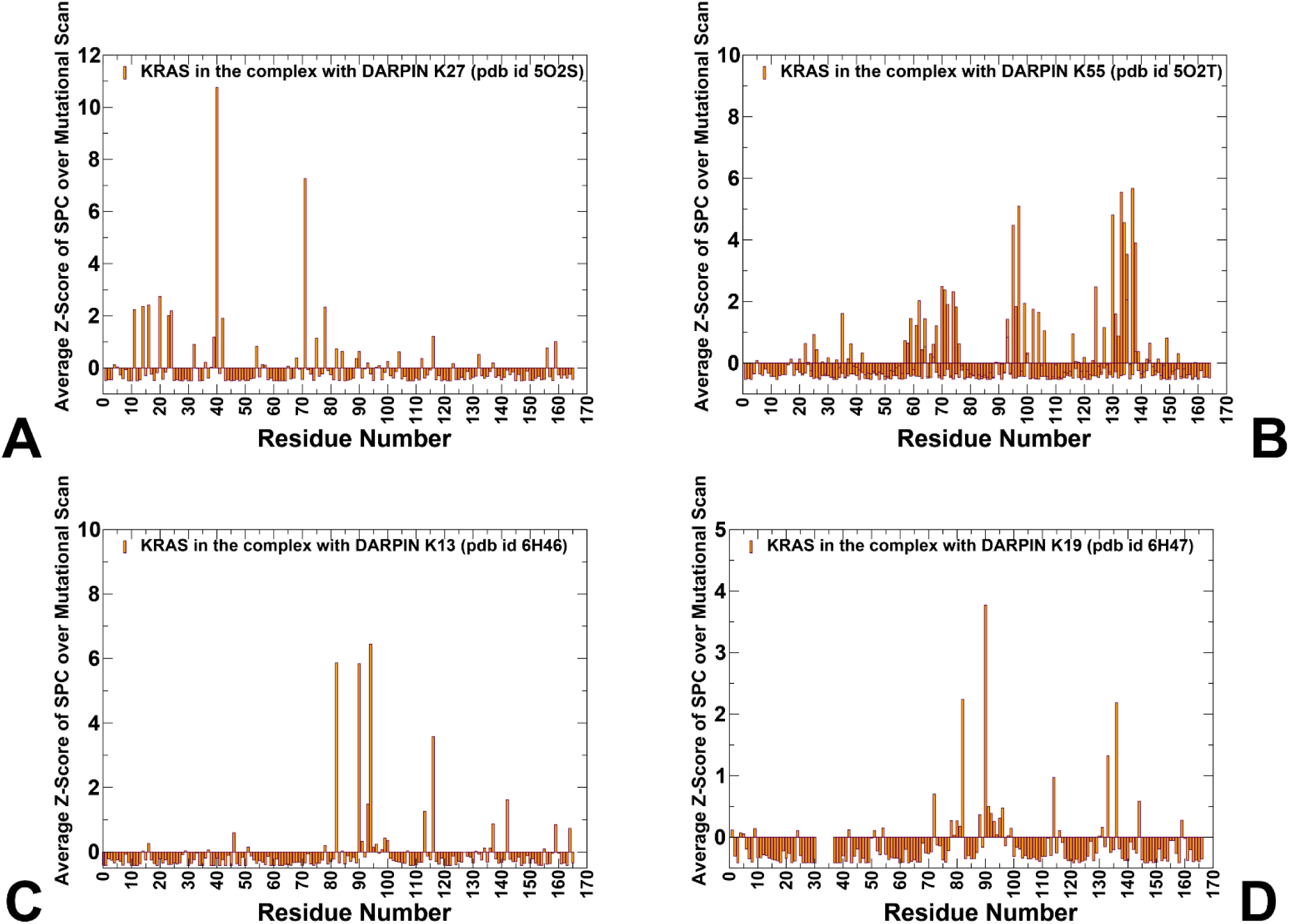
Network centrality analysis of the KRAS complexes with DARPIN proteins. The average Z-score of the ASPL over mutational scan is obtained for each node based on the change of the characteristic path length under node removal averaged over all protein residues. The Z-score are then computed over all mutational changes in given position. The network metric profile corresponding to the average Z-score of the ASPL over mutational scan for the KRAS residues in the KRAS-WT complex with DARPIN K27 (pdb id 5O2S) (A), KRAS-WT complex with DARPIN K55 (B), KRAS-WT complex with DARPIN K13 (C) and KRAS-WT complex with DARPIN K19 (D). The profiles are shown in orange-colored filled bars.

Despite both targeting the effector lobe, the Z-score profiles show that K27 engages a broader, distributed network where mutations at structurally distant residues can severely disrupt long-range signaling, even when they are not directly involved in binding. K55 focuses on a more compact, effector-like topology, where disruption of key Switch II and allosteric lobe motifs has disproportionate effects on network efficiency, indicating a higher sensitivity to aromatic loss and local destabilization.

Unlike K27, Z-score RCA peaks for KRAS residues in the complex with K55 are concentrated around Switch II and the allosteric lobe, suggesting that it relies on localized signal reinforcement rather than broad network modulation. The fact that mutations at Y137, L133, and R97 significantly impair network efficiency indicates that K55 exploits not only the effector interface, but also regions previously thought to be functionally remote, such as Loop 7 and helix α4. For K13, the major mutation-sensitive cont5rol points are M78, F90, H94, F82, L113, N116, I142, Y137, L159 (Figure 8C). For K19 the hubs are generally similar but not identical : F90, F82, S136, L133, V114, M72, V114, T144 (Figure 8D). The important finding of this analysis is that the predicted allosteric hotspots are consistent with the seminal experimental studies of KRAS allostery showing that allosteric effects in KRAS are mediated by the central β-strand of KRAS which acts as a hub for transmitting conformational changes linking distant functional sites [34]. According to DMS analysis of KRAS allostery KRAS positions F78, F82 from β-strand 4 and residues V112, V114 and N116 from β-strand 5 can act as allosteric centers affecting binding with effector proteins through long-range interactions. The important revelation of our predictions is that the allosteric binding hotspots correspond to conserved positions that could control allosteric binding with diverse range of binding partners including DARPin proteins.

The central revelation of our predictions is that the allosteric binding hotspots correspond to conserved positions that could control allosteric binding with diverse range of binding partners targeting different pockets on KRAS. In addition, a group of residues from the α-helix 3 (I93, H95 and Y96) is critical for linking the binding interface region with the central β-strand and subsequently to the allosteric lobe. Finally, the predicted allosterically important positions (Y157, L159, I163 and R164 from α-helix 5) were found in the experimentally validated allosteric pocket located in the C-terminal lobe of the protein (this pocket is formed by residues 97, 101, 107–111, 136–140 and 161–166)and is the most distant pocket from the binding interface [34]. Our analysis identifies these sites as major hotspots of allosteric communication that affect interaction paths from remote parts of KRAS to the binding interface.

## Discussion

KRAS has long stood as a paradigm of functional complexity—its conformational transitions between GDP- and GTP-bound states, its reliance on switch regions for effector engagement, and its integration into signaling networks have made it both an essential regulator and a formidable therapeutic target. The emergence of engineered binders like DARPINs K27, K55, K13, and K19 has opened new avenues for targeting this oncogenic hub with precision and selectivity. These systems do not simply block KRAS function; they engage it through diverse molecular strategies that span effector mimicry, conformational restriction, and allosteric modulation. By combining mutational scanning, MM-GBSA decomposition, RMSF profiling, and network-based analysis, we uncover a layered understanding of how these DARPINs exploit the structural and energetic architecture of KRAS to modulate its activity. At the heart of our findings is the realization that KRAS recognition by engineered proteins is not monolithic. Instead, it reflects a spectrum of binding mechanisms, each with unique implications for mutation sensitivity and functional interference. The comparison between K27 and K55 reveals a striking dichotomy in the way engineered proteins can engage KRAS. While both target the classical effector interface involving Switch I and II loops, their mechanisms diverge significantly. K27 binds to the inactive, GDP-bound state, stabilizing the nucleotide-binding pocket without inducing major structural rearrangements. Key residues such as R41, T35, and Y40 contribute through modular hydrogen bonding and electrostatic anchoring, forming a broad interaction scaffold rather than relying on localized high-affinity contacts. In contrast, K55 mimics natural effectors like RAF1 and PI3Kγ, locking KRAS into a signaling-ready geometry through a compact hydrophobic cluster centered on Y40, M67, and F79, supported by electrostatic interactions at D38 and R41. This system acts more like a molecular trap, interfering with downstream signal propagation by restricting motion in both switch regions and altering the electrostatic environment around the effector lobe.

Mutational scanning offers a powerful lens through which to evaluate residue-level contributions to complex stability, revealing a core set of evolutionarily constrained hotspots that are critical for recognition across all systems. Notably, I36 and Y40 emerge as universal hubs, contributing significantly to binding in both natural effectors and engineered proteins. However, mutational effects diverge between DARPINs. For instance, R41 becomes the dominant hotspot in K27, where mutations such as R41C/H/N/D cause severe destabilization due to loss of salt bridge formation and backbone-mediated hydrogen bonding. By contrast, Y64 and M67 are uniquely vulnerable in K55, with substitutions like Y64G or M67L/V/A leading to dramatic reductions in binding affinity—likely due to disruption of aromatic stacking and shape complementarity. These results underscore that while certain residues serve as central anchors across multiple binders, others define specificity and escape vulnerability, highlighting the importance of tailoring therapeutic strategies to match the energetic and mutational context of each system.

The MM-GBSA residue-wise decomposition deepens this mechanistic picture, showing that hydrophobic forces dominate interface stabilization, even when electrostatic contributions vary significantly between systems. In K27, the electrostatic environment around R41 plays a defining role, with a total contribution of ΔG = -16.34 kcal/mol, driven primarily by favorable polar interactions. This contrasts sharply with K55, where van der Waals contacts at Y40 and M67 form the backbone of binding stability. Despite their shared engagement of the effector lobe, the energetic dependency of these two DARPINs reflects their distinct functional roles: K27 acts as a conformational restrainer, preventing SOS1-mediated activation, while K55 functions as an effector mimic, blocking downstream signal propagation by locking KRAS into a rigid, signaling-ready orientation. This divergence in thermodynamic drivers provides a rationale for combining these agents in therapeutic regimens, as their overlapping yet mechanistically distinct modes of action raise the genetic barrier to resistance. Beyond the classical effector interface, K13 and K19 offered a different strategy in KRAS targeting, engaging a structurally remote allosteric lobe centered on the α3–loop–α4 motif. This pocket includes H95, Y96, L133, S136, and Y137, with D108 emerging as a defining feature of K19, where it forms salt bridges and hydrogen bonds that enhance selectivity for KRAS over HRAS and NRAS. Unlike K27 and K55, K13 and K19 do not fully restrict switch loop motion, but instead alter the conformational equilibrium of KRAS, subtly shifting the balance toward functionally constrained states. This dynamic modulation aligns with experimental observations that these DARPINs allow for partial nucleotide exchange and effector engagement, distinguishing them from strict pathway blockers and positioning them as fine-tuners of KRAS activity rather than complete suppressors.

Network-based analysis reveals that the impact of DARPin binding extends beyond the immediate interface. High Z-score peaks in K27-bound KRAS include Y40, Y71, T20, K16, V14, L23, F78, S39, and N116, many of which are not directly involved in binding but function as bottlenecks in long-range communication. These residues contribute to maintaining the structural integrity of the central β-sheet, suggesting that K27 indirectly reshapes the global network topology, favoring inactive-state communication routes. In contrast, K55 shows elevated Z-scores at Switch II residues such as Q70, Y71, T74, Y137, L133, R97, A130, and A134, indicating that its mechanism relies on localized rigidity and enhanced effector-like connectivity. Mutations at these positions severely disrupt network efficiency, supporting the idea that K55 reinforces the active-state residue interaction map, making it overly sensitive to aromatic loss but less affected by upstream P-loop variations. The comprehensive analysis of KRAS binding with engineered DARPINs—K27, K55, K13, and K19—reveals a strikingly consistent theme: despite their distinct mechanisms of recognition, all systems engage a unifying allosteric architecture that spans multiple functional motifs. This architecture is not only preserved across complexes but also mirrors the intrinsic communication framework of KRAS itself, where specific residues function as central hubs, transmitting conformational changes across the protein. Notably, this includes positions such as H94, H95, Y96, L133, S136, Y137, and D108 (in K19) —residues located within the allosteric lobe formed by helix α3–loop–α4, which has been experimentally validated as a key regulatory domain for modulating RAS function.

This study reveals that KRAS binding with DARPINs and small molecule inhibitors follows a common architectural logic, governed by a conserved network of residues that span both the effector interface and the allosteric lobe. The Z-score RCA and SPC profiles highlight how mutations at seemingly remote sites—such as N116, F78, and H95 —can exert profound effects on binding stability and signaling output, reinforcing the idea that KRAS functions as an integrated communication system, where local perturbations reshape global dynamics. Moreover, the overlap between allosteric centers recognized by DARPINs and those targeted by small molecule inhibitors like MTX-1106 and Sotorasib [46] suggests that this remote lobe is not only structurally conserved, but also evolutionarily privileged for therapeutic intervention. The presence of H95 and Y96 as central hubs further supports the notion that KRAS-specific allostery can be harnessed for precision oncology. Ultimately, our findings suggest that the future of KRAS inhibition lies in exploiting this universal architecture, where modular hydrogen bonding, aromatic stacking, and network-level restructuring combine to deliver potent, durable, and mutation-resilient targeting of one of cancer’s most persistent drivers.

## Conclusions

This study presents a comprehensive and integrative computational analysis of KRAS binding with engineered DARPINs, combining conformational dynamics, mutational scanning, MM-GBSA residue decomposition, and network-based centrality profiling to uncover the molecular mechanisms that define binding specificity, mutation sensitivity, and allosteric control. These findings provide a detailed mechanistic framework for understanding how distinct classes of binders—effector mimics, conformational blockers, and allosteric modulators—exploit both local and global features of KRAS to achieve targeted inhibition. MD simulations reveal that KRAS exhibits significant conformational plasticity, even under bound conditions, with key functional regions such as the P-loop, Switch I, and Switch II displaying distinct flexibility patterns depending on the nature of the interaction partner. Notably, K27 and K55 stabilize the effector lobe through different dynamic strategies. K27 restricts P-loop, and Switch I motion, locking KRAS into an inactive-state conformation without inducing large-scale structural rearrangements. K55 immobilizes both switch loops simultaneously, mimicking natural effectors like RAF1.These results highlight that functional outcomes are not solely determined by static structures, but rather by how each binder reshapes the intrinsic flexibility landscape of KRAS.

Mutational scanning across all four DARPIN systems identifies a core set of evolutionarily constrained residues that function as universal hotspots in KRAS recognition. KRAS residues I36, Y40, M67, and H95 consistently emerge as critical contributors to binding stability. Importantly, mutational effects extend beyond direct contact points. Residues not directly involved in binding such as R41, F78, and N116 also show high sensitivity to mutation, indicating their role in supporting long-range communication pathways and modulating network efficiency. This emphasizes that KRAS function is governed by distributed interactions, where mutations at structurally distant sites can have profound consequences on signaling output. The MM-GBSA residue-based energy decomposition provides a quantitative view of these interactions, showing that K27 relies heavily on electrostatic contributions, especially from R41, which forms salt bridges and hydrogen bonds that anchor the interface. K55 exploits a dense hydrophobic cluster involving Y40, M67, and F79, with electrostatic networks at D38 and E37 enhancing its effector-mimetic signature. K13 and K19 engage a novel allosteric pocket centered on helix α3–loop–α4, where H95, Y96, L133, and D108 (in K19) contribute strongly to binding stability via aromatic stacking and polar interactions. These energetic profiles confirm that while hydrophobic forces dominate interface stabilization, electrostatic contributions play a defining role in determining specificity.

Network-based analysis reveals that DARPin binding restructures the global communication architecture of KRAS, even when engaging structurally remote pockets. We found that KRAS allostery involves not only localized conformational changes but also global network restructuring, reinforcing the idea that effective targeting requires consideration of both direct and indirect effects on protein communication. This work establishes that KRAS operates within a highly coordinated communication infrastructure, where certain residues function as central hubs governing both local binding and global conformational transitions. What emerges from our findings is a universal network topology underlying KRAS allostery, where certain positions are conserved across diverse binders —both natural effectors and synthetic inhibitors—and play critical roles in shaping the energetic, dynamic, and network-level properties of the protein. This suggests that KRAS does not simply fold into isolated functional units; rather, it operates as an integrated signaling module, with evolutionarily constrained hotspots that govern both local binding events and long-range signal propagation. Notably, several of these hotspot residues— including H94, H95, Y96, are engaged by both small molecule inhibitors and protein-based binders suggesting universally conserved allosteric architecture. This convergence across modalities underscores the evolutionary importance of this region and supports its use as a therapeutic nexus for isoform-specific modulation. By integrating dynamic, energetic, and network-level insights, we demonstrate that effective KRAS targeting must move beyond traditional views of local binding motifs and instead embrace a systems-level approach, where residue interaction networks and hinge point redistribution guide the design of next-generation inhibitors.

## Supplementary Materials

The following supporting information can be downloaded at: www.mdpi.com/xxx/s1, Figure S1 presents structural mapping of conformational mobility profiles of KRAS-DARPin complexes obtained from MD simulations. Figure S2 presents structural mapping of allosteric communication hotspots in KRAS complexes with DARPin proteins.

## Author Contributions

Conceptualization, G.V.; Methodology, M.A., V.P., B.F., G.V.; Software, M.A., V.P., B.F., G.V.; Validation, G.V.; Formal analysis, G.V., M.A., V.P.,B.F.; Investigation, G.V. ; Resources, V.P., M.A, G.V. ; Data curation, M.A., V.P., G.V.; Writing— original draft preparation, G.V.; Writing—review and editing, G.V.; Visualization, V.P., M.A., G.V. Supervision G.V. Project administration, G.V.; Funding acquisition, G.V. All authors have read and agreed to the published version of the manuscript.

## Institutional Review Board Statement

Not applicable.

## Informed Consent Statement

Not applicable.

## Data Availability Statement

The original contributions presented in this study are included in the article/supplementary material. Crystal structures were obtained and downloaded from the Protein Data Bank (http://www.rcsb.org). The rendering of protein structures was done with UCSF ChimeraX package (https://www.rbvi.ucsf.edu/chimerax/) and Pymol (https://pymol.org/2/). All mutational heatmaps were produced using the developed software that is freely available at https://alshahrani.shinyapps.io/HeatMapViewerApp/.

## Acknowledgments

The authors acknowledge support from Schmid College of Science and Technology at Chapman University for providing computing resources at the Keck Center for Science and Engineering.

