## Supplemental Figures S1,S2 for "A Conserved Architecture of Allosteric Communications and Regulatory Hotspots in the KRAS Complexes with Diverse Binding Proteins Controls Mechanisms of Effector Mimicry and Allosteric Modulation : Atomistic Revelations from Molecular Simulations and Mutational Scanning of Binding Energetics and All"

**Supp****lementary Material****s**

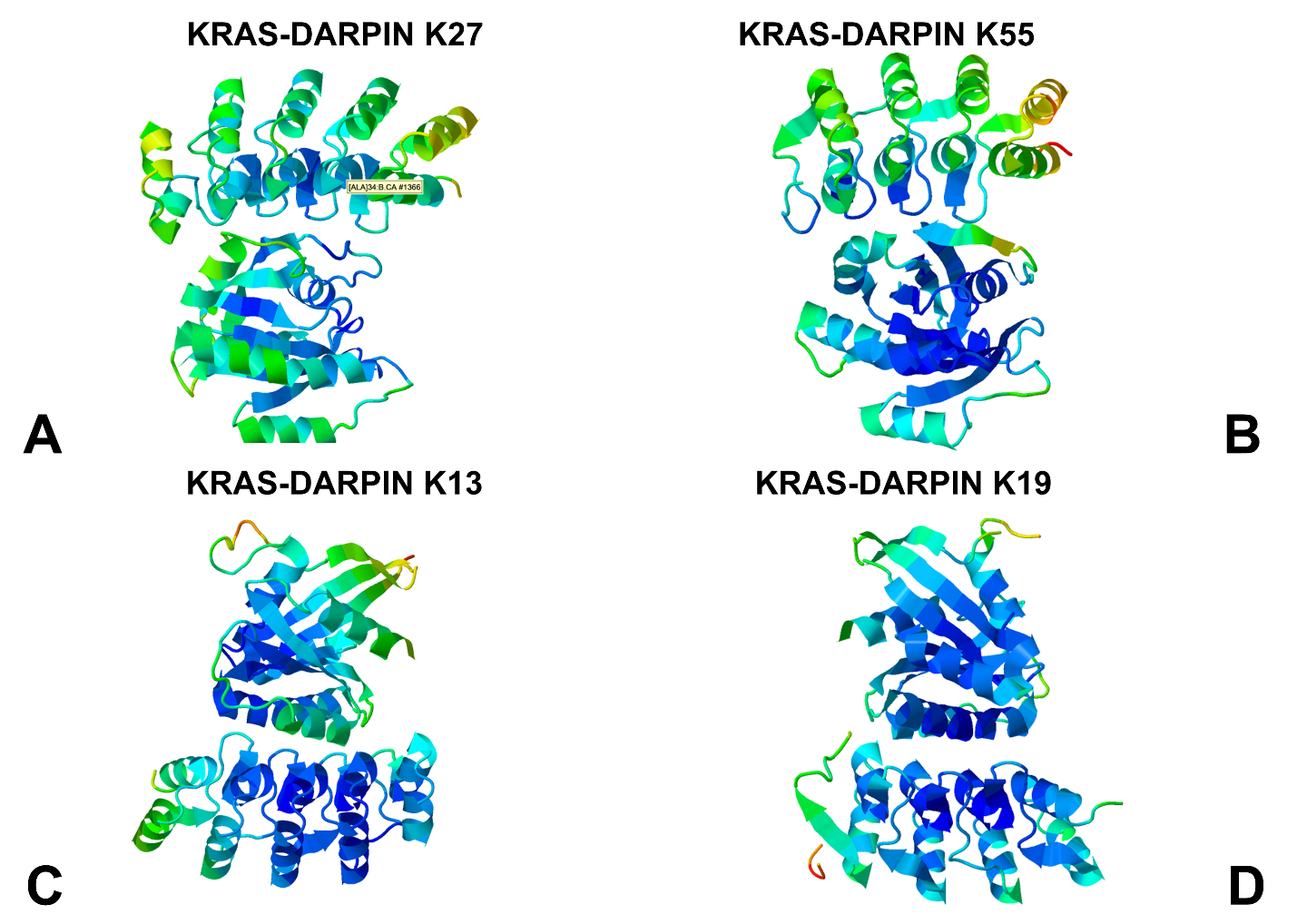

**Figure S1.** Structural mapping of conformational mobility profiles obtained from MD simulations of KRAS complexes with DARPin proteins K27 (A), K55 (B), K13 (C) and K19 (D) . The structures are shown in ribbons with the rigidity-to-flexibility scale colored from blue to red with the most immobilized in functional motions regions in blue and most flexible in red.

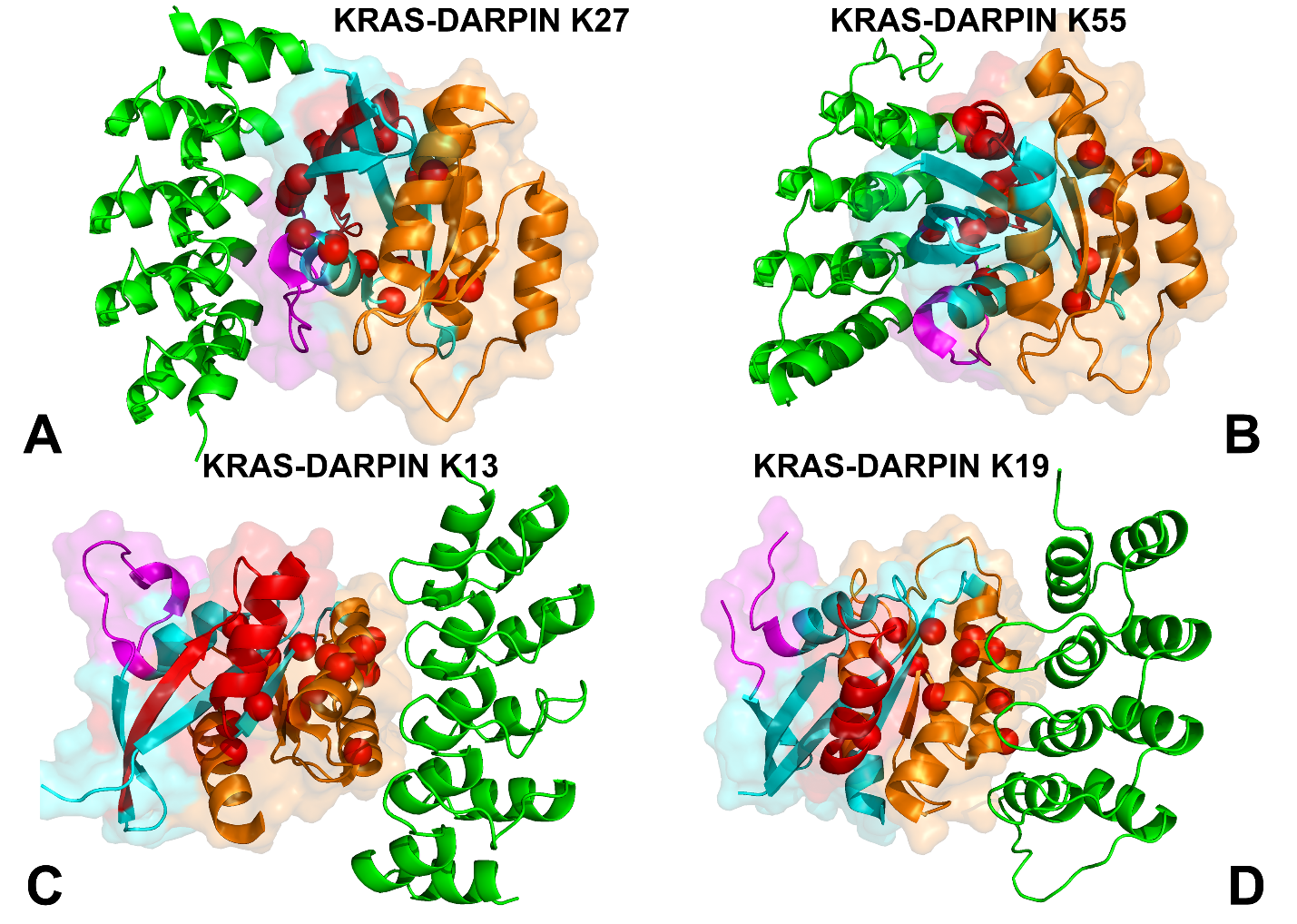

**Figure S2.** Structural mapping of allosteric communication hotspots corresponding to the major peaks of the SPC network centrality profile for KRAS complexes with DARPin proteins K27 (A), K55 (B), K13 (C) and K19 (D) . KRAS protein is shown in ribbons and transparent surface. The switch I region (residues 24-40) is in magenta, switch II region (residues 60-76) is in red and allosteric lobe (residues 87-170) is in orange. The DARPin proteins are shown in green ribbons. The allosteric communication hotpots are shown in red spheres.
